# SAM-dependent viral MTase inhibitors: herbacetin and caffeic acid phenethyl ester, structural insights into dengue MTase

**DOI:** 10.1101/2022.05.31.494145

**Authors:** Mandar Bhutkar, Amith Kumar, Ruchi Rani, Vishakha Singh, Akashjyoti Pathak, Aditi Kothiala, Supreeti Mahajan, Bhairavnath Waghmode, Ravi Kumar, Rajat Mudgal, Debabrata Sircar, Pravindra Kumar, Shailly Tomar

**Author notes:** Corresponding author address: Department of Biosciences and Bioengineering, Indian Institute of Technology Roorkee, Roorkee-247667, Uttarakhand, India; Email id.

## Abstract

Chikungunya (CHIKV) and dengue (DENV) viruses pose a public health risk and lack antiviral treatment. Structure-based virtual screening of natural MTase substrates library identified herbacetin (HC) and caffeic acid phenethyl ester (CAPE) as potential CHIKV nsP1 and DENV NS5 MTase inhibitors. Binding affinities and MTase inhibition were confirmed using purified proteins. Crystal structure of DENV3 NS5 MTase and CAPE complex revealed CAPE binding at GTP and cap 0 RNA sites. Interestingly, HC and CAPE depleted polyamines, which are crucial for RNA virus replication, and effectively diminished replication with IC_50_ values of ∼13.44 µM and ∼0.57 µM against CHIKV, and ∼7.24 µM and ∼1.01 µM against DENV, respectively. Polyamine addition did not reverse the antiviral effects, suggesting a dual inhibition mechanism.

**Graphical abstract:** 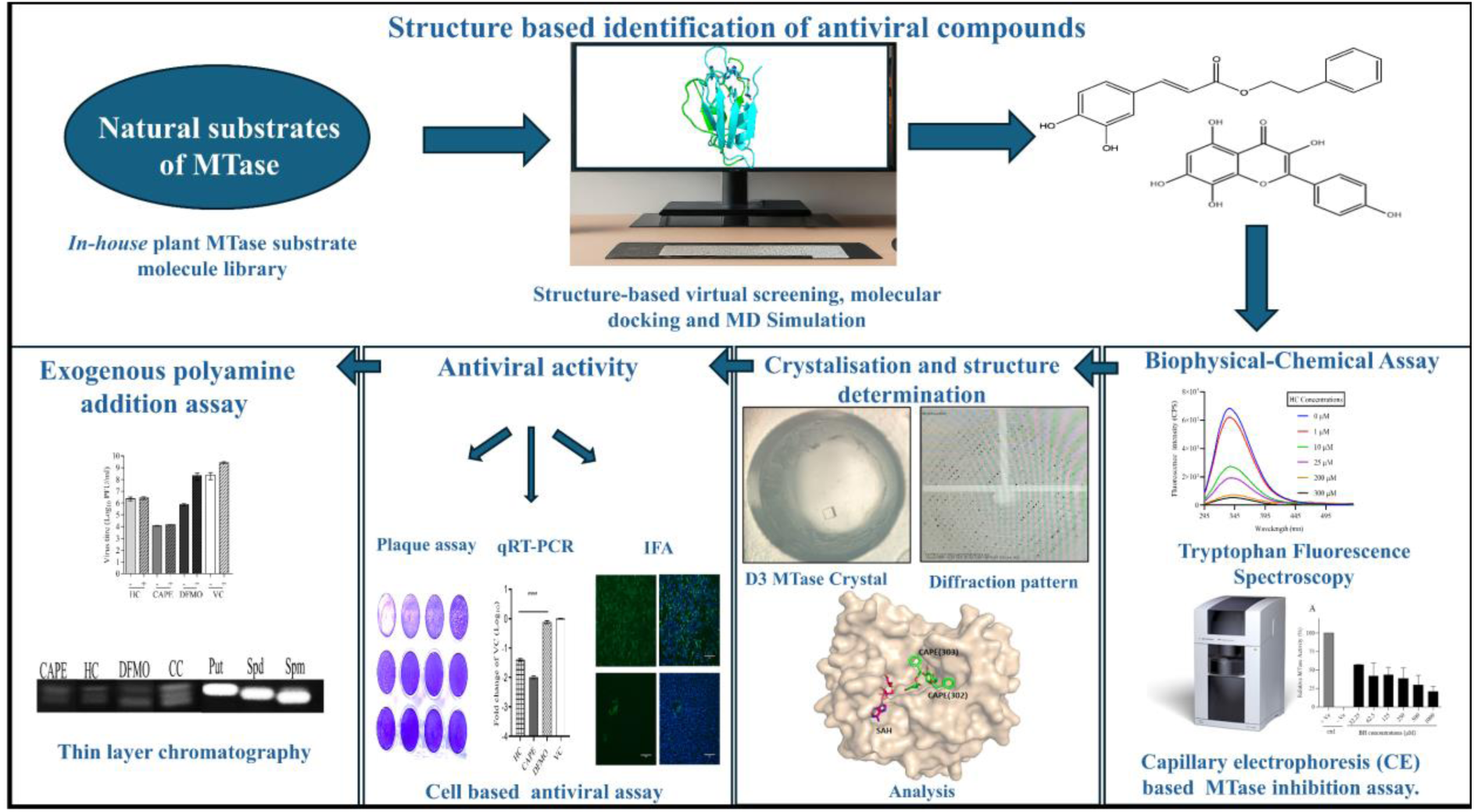

## Introduction

Dengue virus (DENV) and chikungunya virus (CHIKV) are arthropod-borne viruses belonging to the *Flaviviridae* and *Togaviridae* families, respectively (1). Both viruses cause febrile illnesses, showing fever, arthralgia, joint pain, rash, and headache. These enveloped viruses have a positive-sense single-stranded RNA genome (+ssRNA) (2). It is estimated that annually, ∼100-400 million DENV infections occur worldwide. The Philippines and Vietnam are among the most affected countries. Moreover, dengue is endemic in India, Indonesia, Myanmar, Sri Lanka, and Thailand (3–5). Further, in 2023, ∼ 0.5 million CHIKV cases occurred worldwide, of which most were reported in Brazil and India (6,7). In India, co-infections with DENV and CHIKV occur prevalently in various regions during the monsoon season (6,8,9).

The *Togaviridae* family includes various alphaviruses such as CHIKV, Eastern equine encephalitis virus (EEEV), Venezuelan equine encephalitis virus (VEEV), etc. Alphaviruses comprise four nonstructural proteins (nsPs), namely nsP1, nsP2, nsP3, and nsP4, which play an essential role in the life cycle of the viruses and are produced when a viral protease, i.e., nsP2, cleaves the viral polyprotein precursor post-translationally (10). Among the four nonstructural proteins (nsPs) in alphaviruses, nsP1 is crucial due to its essential role in viral RNA capping and is a potential drug target (11–14). The viral RNA caps have numerous biological roles, such as helping the eukaryotic translational initiation factor 4E (eIF4e) recognize the RNA for initiating translation and protecting the mRNA from cellular exonucleases (15). Further, the cap structure also helps the viral genome to escape recognition from the host innate immune system pathways such as retinoic acid-inducible gene I (RIG-I) and melanoma differentiation-associated protein 5 (MDA5) (16). In alphaviruses, the cap 0 structure is characterized by monomethylation at the N7 position of the guanosine nucleotide(17). The capping enzyme nsP1 of alphaviruses facilitates viral RNA capping through two crucial steps. Initially, S-adenosyl methionine (SAM) serves as a methyl donor, transferring a methyl group to the seventh position of GTP, resulting in the formation of m7 guanosine-5’-monophosphate (m7GMP) and producing S-adenosyl homocysteine (SAH) as a by-product. This process is referred to as the MTase step of the capping reaction (18). In the subsequent guanylation (GT) step, the methylated m7GMP is covalently attached to His37 of the nsP1 enzyme and is transferred to the 5’ end of the viral RNA (18).

The *Flaviviridae* family includes orthoflaviviruses like DENV, Zika (ZIKV), Japanese encephalitis, etc. DENV consists of four distinct serotypes: DENV 1, DENV 2, DENV 3, and DENV 4. DENV-2 has been the predominant serotype in India for the past five decades. Nevertheless, recent epidemics have also witnessed the emergence of serotypes 3 and 4, indicating a dynamic shift in the circulating dengue virus serotypes in the region (19). Orthoflaviviruses contain a single open reading frame, which encodes a polyprotein processed by viral and cellular proteases into three structural and seven nonstructural proteins (NS1, NS2A, NS2B, NS3, NS4A, NS4B, and NS5). Among these non-structural proteins, NS5 comprises of an N-terminal methyltransferase (MTase) domain and a C-terminal RNA-dependent RNA polymerase (RdRp) domain, each performing the functions of viral RNA capping and RNA synthesis, respectively (20–22). The mechanism of flaviviral MTase involves a two-step process. In the first step, the enzyme transfers a methyl group from SAM to the guanine base located at the 5’ end of the viral RNA, forming a cap 0 structure. In the second step, this cap 0 structure undergoes further modifications to form the cap 1 structure, achieved by adding a methyl group to the ribose sugar of the initially transcribed nucleotide (23,24).

Additionally, polyamines have been shown to be essential for the viral life cycle in these RNA viruses (25,26). There are three biogenic polyamines present in the host of these viral pathogens, namely, putrescine (put), spermine (spm), and spermidine (spd), which are produced from ornithine through the polyamine biosynthetic pathway (27). Enough evidence suggests that polyamines are essential for various stages of the virus life cycle, including genome replication, virus protein translation, and genome packaging (26). The inhibition of the first enzyme, ornithine decarboxyalse 1 (ODC1) of polyamine biosynthetic pathway has been shown to exhibit antiviral activity against CHIKV, DENV, ZIKV etc (25,26). Difluoromethylornithine (DFMO) is a well-known irreversible ODC inhibitor but is associated with hearing loss and may lead to antiviral resistance because depletion of polyamines in CHIKV infected cells leads to muatations in nsP1 and also in the Opal stop codon (28,29). Additionally, a higher dose of DFMO is required for effective antiviral activity, and DFMO is also known to induce the uptake of putrescine and spermidine (30).

The dodecameric cryo-electron microscopy (Cryo-EM) structures of CHIKV nsP1 (PDB IDs: 6Z0V, 6Z0U, and 7DOP) reveal that the upper ring exhibits bifunctional MTase/GTase catalytic activity, whereas the lower ring is responsible for oligomerization and membrane binding (11,12). Similarly, the dimeric structure of DENV NS5 has been reported (PDB ID: 4V0R), revealing its organization of functional domains. Recently, the crystal structure of the NS5 MTase from the Omsk hemorrhagic fever virus (OMHV), which belongs to the *Flaviviridae* family, has been determined. This structure reveals new insights, including the presence of GMP derived from GTP through the action of guanyl transferase (GTase), GMP-arginine adducts, and an uncommonly observed capped RNA conformation (21).

SAM-dependent MTases are categorized as N-, O-, or C-MTases (N-MTases, O-MTases, C-MTases) according to the specific atom (nitrogen, oxygen, or carbon) they methylate in their substrates (31). These enzymes utilize SAM to transfer methyl groups to compounds such as flavonoids and phenolic acids, altering their chemical properties and biological activities. Given their specificity for such substrates, employing these compounds as potential inhibitors is a logical approach for studying enzyme inhibition (31,32). The structure of DENV NS5 MTase and CHIKV nsP1 belong to a family of SAM-dependent MTases (12,33). Moreover, the “Rossmann fold” is a βαβ super-secondary structure that forms the catalytic core responsible for SAM binding site in all SAM-dependent MTases, including DENV NS5 MTase domain and CHIKV nsP1 protein (34–37). Here, highly conserved acidic residues interact with the ribose moiety of SAM, while the glycine-rich loop region interacts with its adenosine moiety (35).

These structural insights enhance the efficiency of high-throughput computational screening for novel natural compounds. Identifying new antiviral agents is a multifaceted and resource-intensive endeavor. Natural compounds, ubiquitously present in various food items, offer the advantage of potentially exhibiting a lower side effect profile than synthetic pharmaceuticals (38). Consequently, structure-based virtual screening represents a promising approach for identifying compounds with potential pharmacological activity (39). In the current study, an *in house* library of natural substrates of N-C-O-MTases (NSMT), was curated and screened against CHIKV nsP1 and DENV NS5 MTase. High throughput virtual screening and experimental validation pinpointed Herbacetin (HC) and Caffeic acid phenethyl ester (CAPE) as potent MTase inhibitors.

## Materials and methods

### Multiple Sequence Alignment (MSA)

The amino acid sequence of the nsP1 protein of alphaviruses was compared with CHIKV nsP1, and the NS5 MTase domain of orthoflaviviruses was compared with the DENV 3 NS5 MTase protein. CHIKV nsP1 and DENV 3 NS5 MTase were utilized as reference points for these alignments. The sequence alignment profile of the selected nsP1 and NS5 MTase sequences was performed via Clustal Omega tool and analyzed by a graphical colored depiction using ESPript 3.0 (40). A comprehensive description of the protocol is given in the supplementary materials.

### Structure-guided virtual screening of an *in house* library of NSMT

An *in house* NSMT compound library consisting of approximately 25 molecules, comprising flavone and catechol derivatives, was constructed (31). The three-dimensional structures of these molecules were obtained from the PubChem database (https://pubchem.ncbi.nlm.nih.gov/). The library was subsequently applied for virtual screening analyses, employing PyRx 0.8 (41). Each compound was then energy minimized using the Universal Force Field (UFF) in Open Babel and converted to .pdbqt format (42). The Cryo-EM structures of CHIKV nsP1 (PDB ID: 6Z0V) and the crystal structure of DENV-3 NS5 MTase (PDB ID: 4V0R) were retrieved from the RCSB Protein Data Bank (PDB). Given that the structural data for these proteins were incomplete, with several residues in flexible loops missing, SWISS-MODEL was utilized to construct complete models of these proteins, ensuring accurate structural representations for virtual screening (43). Before screening, .PDB were converted into .PDBQT format to facilitate their use in AutoDock Vina for docking (44). The grid focused on the SAM/GTP binding site of CHIKV nsP1 and the GTP binding site of DENV-3 NS5 MTase (41). The GTP site was selected for DENV-3 NS5 MTase due to the frequent presence of SAM/SAH in available crystal structures, indicating a high prevalence of these interactions in the binding site. For CHIKV nsP1, the grid dimensions were set at 33.48 Å × 19.99 Å × 32.10 Å, centered at X = 54.92, Y = 137.80, Z = 91.35, and for DENV-3 NS5 MTase, the grid dimensions were 34.36 Å × 36.68 Å × 24.04 Å, centered at X = 13.79, Y = 134.12, Z = 0.54. Following the initial screening, the top compounds were selected on the basis of binding energy, ligand pose, and key interactions with the target sites as well as the reported antiviral activity, and were docked again using AutoDock Vina (44). The same grid parameters were employed to maintain consistency. The docked conformations were subsequently analyzed using PyMOL for structural visualization and LigPlot+ to comprehensively map hydrogen bonding and hydrophobic interactions (45,46).

### Molecular Dynamics (MD) Simulation

nsP1 apo and in complex with SAM, HC, and CAPE and DENV 3 NS5 MTase apo and in complex with GTP, HC, and CAPE were subjected to MD simulation studies to assess dynamic behavior and stability of protein-ligand interactions. The GROMACS 2022.2 suite was used to carry out all simulation studies using the CHARMM36 force field on a LINUX-based workstation (47,48). Ligand parameters and topology files were generated using the CHARMM General Force Field (CgenFF) program (49). The SPC (Simple Point Charge) water type for solvation and counter ions (Na+ and CL-) were added to neutralize the cubic system. The energy minimization step was performed using the steepest descent method with a maximum force of 10 kJmol-1. Followed by a two-phased equilibration of a constant number of particles, volume, and temperature (NVT), a constant number of particles, pressure, and temperature (NPT) for 100ps each. Periodic boundary conditions were applied at a constant temperature of 300 K and 1 atm pressure, utilizing the V-rescale temperature coupling method and Berendsen pressure coupling method, respectively (50). The long-range electrostatic interactions were calculated using the Particle Mesh Ewald method, while bond lengths involving heavy atoms were constrained with the Linear Constraint Solver (LINCS) algorithm (51). Short-range forces were computed with a minimum cutoff of 1.2 nm, employing the Verlet cutoff scheme (52). Finally, 100 ns MD production run was performed using the leap-frog algorithm with an integration time frame of 2fs, and the trajectories were generated after every 10 ps. The resulting trajectories were analyzed for structural deviations and fluctuations within the protein and protein-ligand complexes .

### Cloning, expression, and purification of DENV 3 NS5 MTase Domain

For the cloning of DENV 3 NS5 MTase Domain, RNA was isolated from the supernatant of DENV 3 infected cells using the TRIzol (Sigma) method described by the manufacturer. The RNA was then reverse-transcribed into complementary DNA (cDNA) using the PrimeScript cDNA Synthesis Kit (Takara). The DENV 3 cDNA was subsequently amplified by PCR, using the following primers (amino acid residues 1-278), NS5 MTase forward, 5′-ATGC<u>GCTAGC</u>GGAACAGGTTCACAAG-3′, and NS5 MTase reverse 5′-CGCA<u>CTCGAG</u>TCAATTAACATGTCGAGTT-3′, the sequences underlined are the recognition sites of the restriction enzymes *NheI* and *XhoI* respectively. After digestion with restriction enzymes, the PCR product was purified with a Qiagen gel extraction kit per the mentioned kit protocol and inserted between the restriction enzymes sites of the vector pET28c (+). The recombinant vector pET28c (+) NS5 MTase was identified by restriction digestion, and the DENV 3-NS5 MTase insert was verified by sequencing. *E. coli* DH5α was used for amplification of the recombinant plasmid, and *E. coli* BL21 (DE3) was transformed for induced expression of the His-tagged DENV 3-NS5 MTase protein. Expression and purification was done, as mentioned in Boonyasuppayakorn et al. 2014 (53). Eluted fractions were dialyzed, and further protein were concentrated to the required concentration with amicon centrifugal filters (10,000 MWCO Millipore, Burlington, MA, USA), flash-frozen in liquid nitrogen, and stored at −80 °C till its use. A detailed description of this protocol is given in the supplementary materials.

### Expression and purification of recombinant CHIKV nsP1

As mentioned by Kaur et al. in 2018, *E. coli* Rosetta cells were utilized for the expression and purification of recombinant CHIKV nsP1 using the recombinant expression plasmid pET28c(+)-CHIKV nsP1 (54,55).

### Tryptophan Fluorescence Spectroscopy

Tryptophan fluorescence spectroscopy (TFS) experiments were conducted using a Fluoromax fluorescence spectrophotometer (Horiba Scientific). A quartz cuvette with dimensions of 5x5 mm was utilized. The excitation wavelength was set to 280 nm, and the emission wavelength was scanned from 295 to 540 nm. A slit width of 5 nm was employed for all measurements. nsP1 and NS5 MTase protein samples were prepared at concentrations of 1 μM and 0.15 μM, respectively, in a 500 μl phosphate-buffered saline (PBS) solution. Variable concentrations of SAM (S-adenosylmethionine), HC, and CAPE were added to the protein samples. The experiments were conducted at 25°C. Control buffer experiments and compound titrations were performed parallel with the main experiments for background determination and used for subtraction. Data from three independent experiments were collected and analyzed using nonlinear regression with the ’One Site-Specific Binding’ model. The data analysis was carried out using GraphPad Prism 8 software (55).

### Capillary electrophoresis (CE) based DENV 3 NS5 MTase and CHIKV nsP1 inhibition assay

To determine the enzyme inhibition activities of HC and CAPE, the enzymatic reaction for of DENV 3 NS5 MTase and CHIKV nsP1 were conducted using specific reaction mixtures. The reaction mixture for NS5 MTase consisted of 50 mM Tris buffer (pH 7.5), 10 mM KCl, 2 mM DTT, 2 mM MgCl_2_, 0.3 mM SAM, and 0.3 mM GTP, along with 1 μM NS5 MTase protein. The same components were used for the nsP1 enzyme reaction with 5 μM nsP1 protein. These enzyme reactions were performed at 37 °C for 1 h. To establish negative controls, both enzyme assays included a reaction with no SAM-GTP (substrate). Furthermore, to provide an additional negative control, the NS5 MTase reaction was also conducted with the inactive capsid of CHIKV. After incubation reaction was stopped by adding acetonitrile in 1:2 (vol/vol), 100 μM caffeine was used as internal control (13,14). The mixture was vortex-mixed for 15 s, and the protein was precipitated for 20 min at 18,500 x g. The supernatant was transferred to sample vials. The detected SAH values were normalized to the internal standard for each capillary electrophoresis run. CE analysis was performed as mentioned in Mudgal et al. 2020. Data from three independent experiments were collected and analyzed using GraphPad Prism 8 software (13,14).

### Crystallization of NS5 MTase

The crystallization of DENV 3 NS5-MTase was conducted using the sitting drop vapor diffusion method in 96-well plates at a temperature of 20°C, as described in Coutard et al., 2014 (56,57). For data collection, DENV 3 NS5-MTase crystals in complex with CAPE were obtained by soaking the crystals in a cryoprotectant solution (22.5% glycerol) containing 1mM CAPE for 1 h. They were flash-frozen in a nitrogen stream at 100K. Diffraction data were collected at the home source (Rigaku Micromax 007 HF), Macromolecular crystallography unit, IIT Roorkee. Data reduction and scaling were done using the CrysAlis Pro software (Rigaku Inc.). The structure was determined by MOLREP using DENV 3 NS5 MTase (PDB code 4R8R) as a search model (58). Iterative rounds of model-building in COOT and the refinement of atomic coordinates and B-factors using refmac5 (59) in CCP4i2 allowed for the correct placement of sidechains and loops. NCS and jelly-body restraints were used throughout the data refinement. Additionally, to confirm ligand localizations in the structure, omit maps were generated in Polder (60). The data collection and refinement statistics are summarised in Table 3. All figures of protein and ligand structures were prepared using PyMOL (45).

### Cell line, Virus isolation, propagation and serotyping

DENV was isolated from the Dengue-suspected patient’s blood samples and was further confirmed as a DENV 3 serotype after sequencing. Vero cells were used for the propagation and titration of DENV and CHIKV. CHIKV (Accession No. KY057363.1.) was propagated and titrated using the protocol reported by Singh et al., 2018 and then stored at −80°C for further experiments (61). A comprehensive description of the protocol is given in the supplementary materials.

### Preparation of HC, CAPE, and DFMO stock solutions

HC, CAPE, and DFMO were purchased from Cayman, USA. For experiments, the stock solutions (HC: 165.45mM; CAPE: 20mM) for both compounds were prepared in dimethyl sulfoxide (DMSO) (Sigma-Aldrich) and filtered through a 0.2 μm size syringe filter (Millipore). The 100 mM DFMO dilution was prepared in sterile tissue culture-grade water that was used as a positive control for the polyamine-related experiments. Further dilutions were prepared in 2% DMEM media (maintenance media) before use. Similarly, stock solutions of individual polyamines (putrescine, spermine, and spermidine) (Alfa-Aesar, USA) were diluted in sterile tissue culture-grade water and used as specified.

### Assessment of Antiviral Activity

Vero cells were seeded onto a 24-well plate at a cell density of 1.0 x 10^5^ cells/well. Compounds with concentrations that maintained cell viability above 90% were used for the treatment. Cells were treated with compounds 12 h before infection. With a multiplicity of infection (MOI) of 0.1, the cell monolayer was infected with the virus with gentle shaking every 15 min for 2 h. The inoculum was removed, and the cell monolayer was washed twice to ensure no chance of secondary infection. Compounds were added in the post-infection (pi) 2% DMEM media and incubated for 24 h. After 24 hpi, the supernatant was collected for the CHIKV antiviral experiment. On the other hand, fresh maintenance media was added to the DENV antiviral experiment for 4 days. Then, the supernatant was collected to determine the virus titer via plaque-forming assay (14,62). A comprehensive description of the cell viability assay and plaque-forming assay given in the supplementary materials.

### Immunofluorescence Assay

Vero cells were seeded onto a 6-well plate at a cell density of 1 x 10^6 cells/well. Cells were treated with compounds, as mentioned earlier. The cells were washed three times with PBS and then fixed with 3.7% formaldehyde for 30 min at room temperature, followed by permeabilization with 0.1% Triton-X-100 for 10 min. After washing the cells with 1X PBS, they were incubated with antibodies against CHIKV and DENV (anti-alphavirus 1:100, Santa Cruz Biotechnology Inc.; 1:500 diluted orthoflavivirus group antibody, Genetex Inc.) for 1 h, followed by a wash with 0.1% Tween-20 in 1X PBS (PBST). Later, the plate was incubated in the dark with fluorescein (FITC)-conjugated secondary anti-mouse antibody (1:250, Sigma) for 30 min at 37 °C. The cells were then washed with PBST and counter-stained with 4′,6-diamidino-2-phenylindole (DAPI, Sigma) for 15 min in the dark. Finally, the images were captured using a fluorescence microscope (EVOS FL AUTO, Thermo Fisher).

### Quantitative Real-Time PCR (qRT-PCR)

HC and CAPE compounds treatment and infection were the same as described in the assessment of antiviral activity. After the termination of an assay, TRIzol was added to the plate. RNA was purified according to the manufacturer’s protocol. Purified RNA was quantified for cDNA preparation using the PrimeScript 1st strand cDNA Synthesis Kit (Takara) with 400 ng of extracted RNA. The KAPA SYBR Fast Universal qPCR Kit was used for qRT-PCR with analysis performed on the QuantStudio™ 5 System from Applied Biosystems, USA. The forward and reverse primers used for amplification are used as previously described in (63) and (55,64). β-actin was utilised as an internal control. The relative quantification was carried out using the ΔΔCt method as described (14,55).

### Polyamine addition assay

In the polyamine addition assay, compounds pre-treatment was given for 24 h at various concentrations such as HC (200 µM), CAPE (25 µM for CHIKV and 2.5 µM for DENV), DFMO (1000 µM). Immediately after 2 h of virus infection with CHIKV and DENV, cells were treated with 1 μM biogenic polyamines (put, spd, spm) for 24 h and 120 h, respectively. Later, the supernatant was collected to determine the virus titer by plaque-forming assay.

### Polyamine determination by Thin-layer chromatography (TLC)

Vero cells were treated with CAPE (25 µM), HC (200 µM), and DFMO (1000 µM) for 36 h. Further, cells were trypsinized, sonicated, and centrifuged. Polyamines were separated by TLC as previously described (65). Briefly, cells were sonicated in 500 µL of 2% (V/V) perchloric acid. Sonication was performed at 4 °C (20 kHz, 2Ω, 10-s pulse, 30-s rest). Cell homogenates obtained are then stored at 4 °C for 24 h. After this, samples are centrifuged at ∼11,500 × g for 30 min at 4 °C. An equal ratio of supernatant dansyl chloride (5 mg/mL) (Alfa-Aesar, USA) was added in acetone and saturated sodium bicarbonate. Samples were incubated in the dark overnight at room temperature. Excess dansyl chloride was cleared by incubating the reaction with 150 mg/mL proline (HiMedia). Dansylated polyamines were extracted with 100 μL toluene. 10 μL of the sample was added in small spots to the TLC plate by automated sprayer (Silica gel matrix; Millipore) and exposed to ascending chromatography with cyclohexane: ethyl acetate (2:3). The plate was dried and visualized via exposure to UV. Further, TLC images were quantified utilizing ImageJ software.

### Statistical analysis

GraphPad Prism 8 software is used for data analysis. One-way ANOVA was used to determine statistical significance wherever mentioned.

## Results

### Structural and sequence conservation of nsP1 and NS5 MTase active sites

Both the viral MTases, nsP1 of CHIKV and NS5 of DENV, bind and use the cofactor SAM as the methyl donor with the release of the byproduct SAH. Structural studies have revealed that the MTase domain in both viral proteins shares common structural features of the Rossmann fold, indicating that these shared characteristics can be exploited to identify a common inhibitor. The SAM binding site in CHIKV nsP1 as revealed by its Cryo EM structure (PDB ID 8AOX, 8AOV) is lined by residues Gly65, Ser66, Ala67, Pro83, Arg85, Ser86, Asp89, Thr137, Asp138, and Gln151. GTP binding site is lined by residues Ala40, Arg41, Ser44, Tyr154, Phe178, Phe241, Val243, Thr246, Tyr248 and Glu250. The residues that make molecular contacts with both SAM and GTP are Arg70, Arg92, and Asp152 (11,12). Similarly, the crystal structural analysis of DENV 3 NS5 MTase (PDB ID 4V0R) revealed that the GTP binding site residues for orthoflaviviruses are Asn15, Arg19, Lys26, Ser147, Arg208, Ser210, Leu14, Phe22 and Thr211 (66,67).

Alphaviruses that are known to cause infections in humans are VEEV, CHIKV, Ross River virus (RRV), Sindbis virus (SINV), Aura Virus (AURA), Middelburg virus (MIDV), Barmah Forest virus (BFV), Madariaga virus (MADV), and Mayaro virus (MAYV). Additionally, Salmonid alphavirus (SAV) exhibits tropism for Atlantic salmon, inducing pancreatic disease, while Eilat virus (EILV) demonstrates insect-specific virulence (68,69). DENV 3, ZIKV, West Nile Virus (WNV), Yellow Fever Virus (YFV), and Japanese Encephalitis Virus (JEV) are some of the human-infecting orthoflaviviruses. In addition, Palm Creek virus (PCV) exhibits insect specificity, and Wenzhou shark flavivirus (WSF) falls within the category of aquatic orthoflaviviruses (70,71). The primary sequence alignment of MTases and subsequent comparison among alphaviruses and orthoflaviviruses demonstrated a significant degree of sequence similarity, as illustrated in supplementary figure 1A. In alphaviruses, 81% of SAM/GTP binding residues in nsP1 are conserved across different viruses. Similarly, in orthoflaviviruses, 66% of GTP binding residues are conserved across NS5 MTase (supplementary figure 1B).

### Structure-based identification of nsP1 and NS5 MTase inhibitors

The generated 3D models of CHIKV nsP1 and DENV 3 NS5 MTase domains showed high structural similarity to templates with root mean square deviation (RMSD) values 0.205 Å and 0.183 Å, respectively (supplementary figure 2 A,B). These models were used for virtual screening of *in house* NSMT library, identifying HC and CAPE as potential inhibitors of nsP1 and NS5 MTase. For nsP1, HC exhibited the highest binding affinity with a binding energy of -8.0 kcal/mol, facilitated by hydrogen bonds with Arg92, Arg70, and Glu88, along with hydrophobic interactions involving Arg41, Tyr285, Asp152, Phe178, Val243, Ala40, and Tyr248 residues nsP1 (Figure 1C, 2C) (Table 1). CAPE demonstrated binding energy of -7.6 kcal/mol, forming hydrogen bonds with Glu250 and Arg92 and engaging in hydrophobic interactions with Tyr248, Arg70, Asp152, Tyr154, Ala40, Phe178, Val243, and Tyr285 residues of nsP1 (Figure 1B, 2B) (Table 1). HC and CAPE exhibited binding affinities comparable to binding affinities of SAM and GTP (positive control) with nsP1 (Figure 1A,D, 2A,D) (Table 1). Similarly, in the case of NS5 MTase, HC displayed binding energy -6.7 kcal/mol with hydrogen bonds formed by Lys11, Asn15, Ser147, and Ser210, along with hydrophobic interactions involving Leu14, Phe22, Pro149, and Thr211 residue of NS5 MTase (Figure 1G, 2G) (Table 1). CAPE exhibited binding energy -6.6 kcal/mol with hydrogen bonds established with Lys11, Leu14, and Leu17 and engaged in hydrophobic interactions with residues such as Asn15, Ser18, and Ser210 residue of NS5 MTase (Figure 1F, 2F) (Table 1). For NS5 MTase, HC and CAPE exhibited slightly lower binding energy than GTP (positive control) (Figure 1E, 2E) (Table 1).

**Figure 1:**
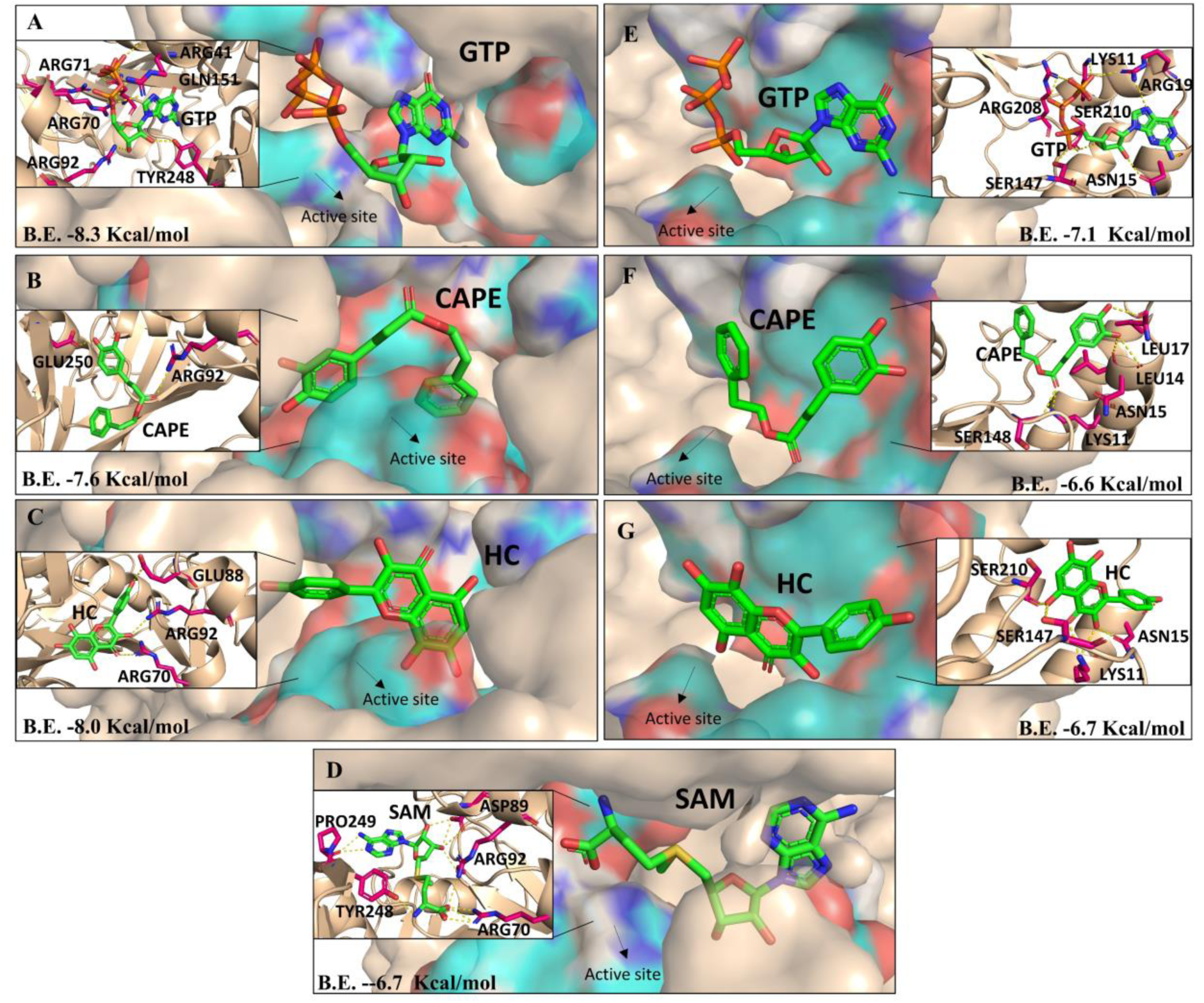
Three-dimensional representation of docked ligands in the enzyme active site (teal surface) of CHIKV nsP1 and DENV 3 NS5 MTase (wheat surface). (A-D) CHIKV nsP1 interacts with the GTP (A), CAPE (B) HC (C), SAM (D), and (E-G) DENV 3 NS5 MTase interact with GTP (E), CAPE (F) HC (G). Zoomed in figures show a detailed view of the binding pocket where molecular interactions of ligands (green) with active site residues (teal colour) and interacting residues (pink colour) of CHIKV nsP1 and DENV 3 NS5 MTase (wheat coloured protein ribbon/ surface).

**Figure 2:**
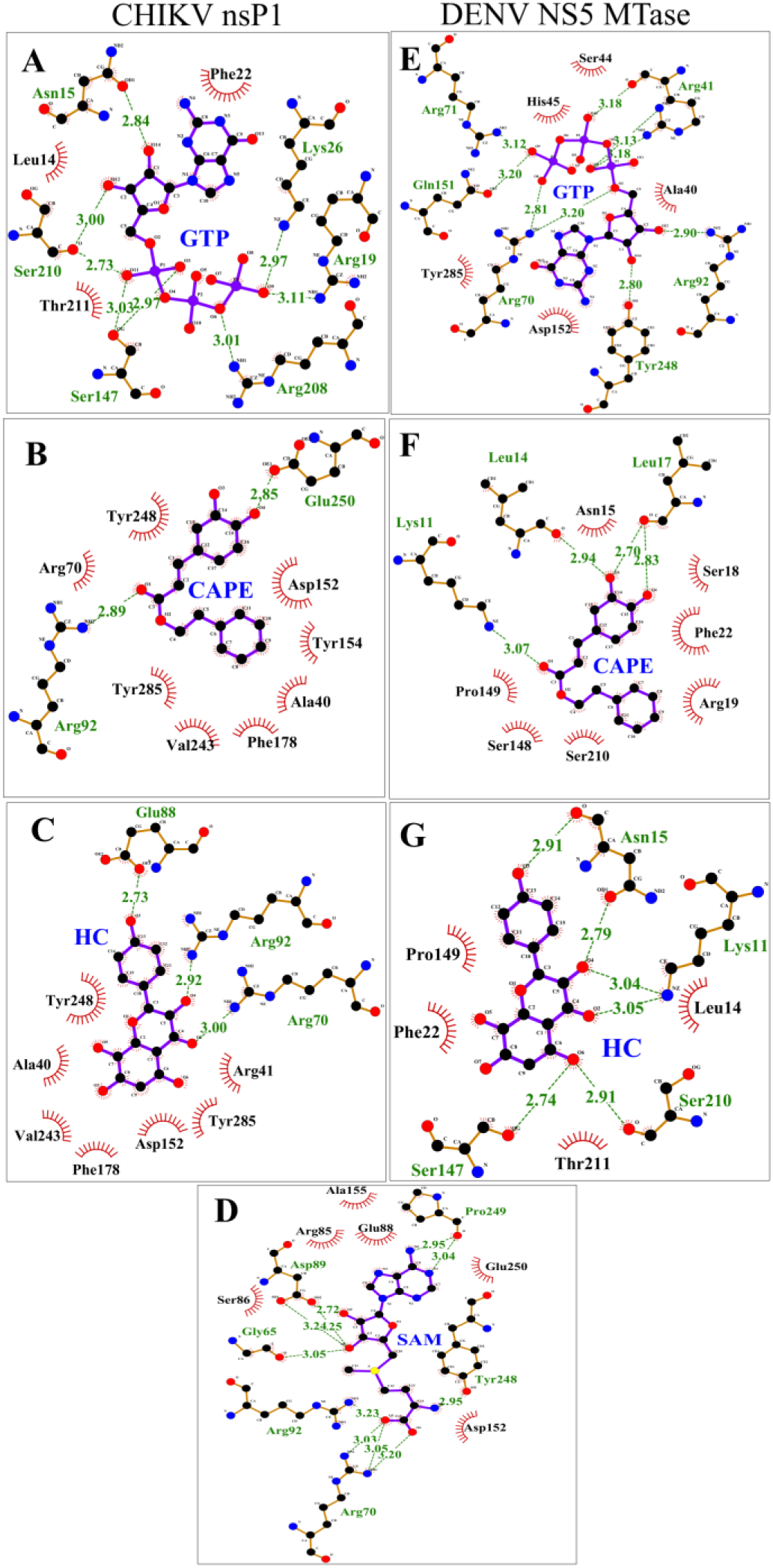
Two-dimensional representation view of docked ligands in the enzyme active site of CHIKV nsP1 and DENV 3 NS5 MTase. (A-D) CHIKV nsP1 interacts with the GTP (A), CAPE (B) HC (C), SAM (D), and E-G) DENV 3 NS5 MTase interact with GTP (E), CAPE (F) HC (G). H-bonds are shown in green dashed lines with the distance shown in Å. Additional residues forming hydrophobic interactions are indicated by a brown semicircle with radiating spokes towards the ligands. 2D interaction figures are made using LigPlot+ software

**Table 1.**
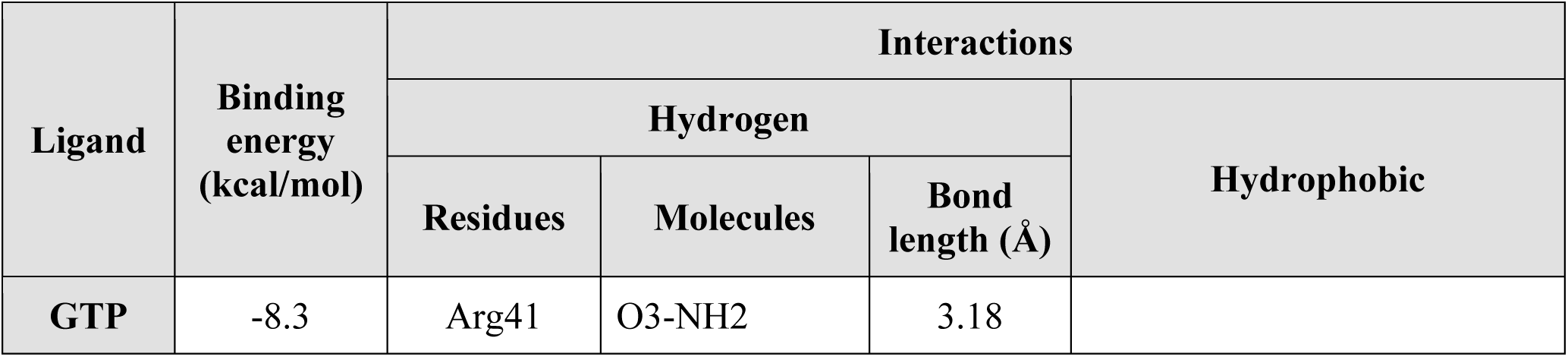
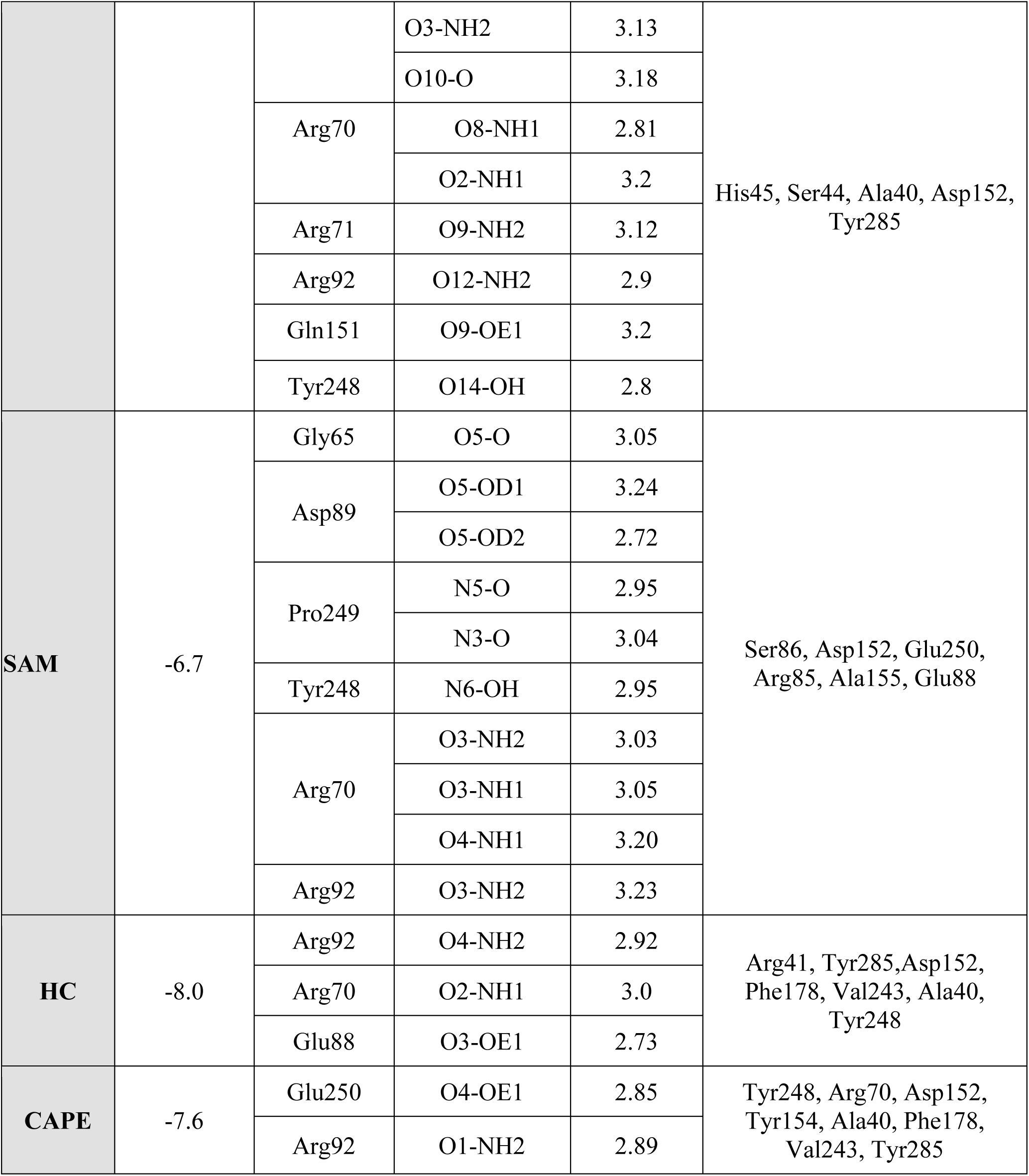
Detailed molecular interactions between the CHIKV nsP1 and ligands.

**Table 2.** Detailed molecular interactions between the DENV 3 NS5 MTase and ligands.

| Ligand |  | Interactions |  |
| --- | --- | --- | --- |
|  |  | Hydrogen | Hydrophobic |

|  | Binding energy (kcal/mol) | Residues | Molecules | Bond length (Å) |  |
| --- | --- | --- | --- | --- | --- |
| **GTP** | -7.1 | Asn15 | O14-OD1 | 2.84 | Leu14, Phe22, Thr211 |
|  |  | Arg19 | O9-NH1 | 3.11 |  |
|  |  | Lys26 | O9-NZ | 2.97 |  |
|  |  | Ser147 | OG-O3 | 2.97 |  |
|  |  |  | OG-O11 | 3.03 |  |
|  |  | Arg208 | O6-NH1 | 3.01 |  |
|  |  | Ser210 | O11-O | 2.73 |  |
|  |  |  | O12-O | 3.00 |  |
| **HC** | -6.7 | Lys11 | O2-NZ | 3.04 | Leu14, Phe22, Pro149, Thr211 |
|  |  |  | O4-NZ | 3.05 |  |
|  |  | Asn15 | O3-O | 2.91 |  |
|  |  |  | O4-OD1 | 2.79 |  |
|  |  | Ser147 | O6-OG | 2.74 |  |
|  |  | Ser210 | O6-O | 2.91 |  |
| **CAPE** | -6.6 | Lys11 | O1-NZ | 3.07 | Asn15, Ser18, Arg19, Phe22, Ser148, Pro149, Ser210 |
|  |  | Leu14 | O3-O | 2.94 |  |
|  |  | Leu17 | O3-O | 2.70 |  |
|  |  |  | O4-O | 2.83 |  |

**Table 3.**
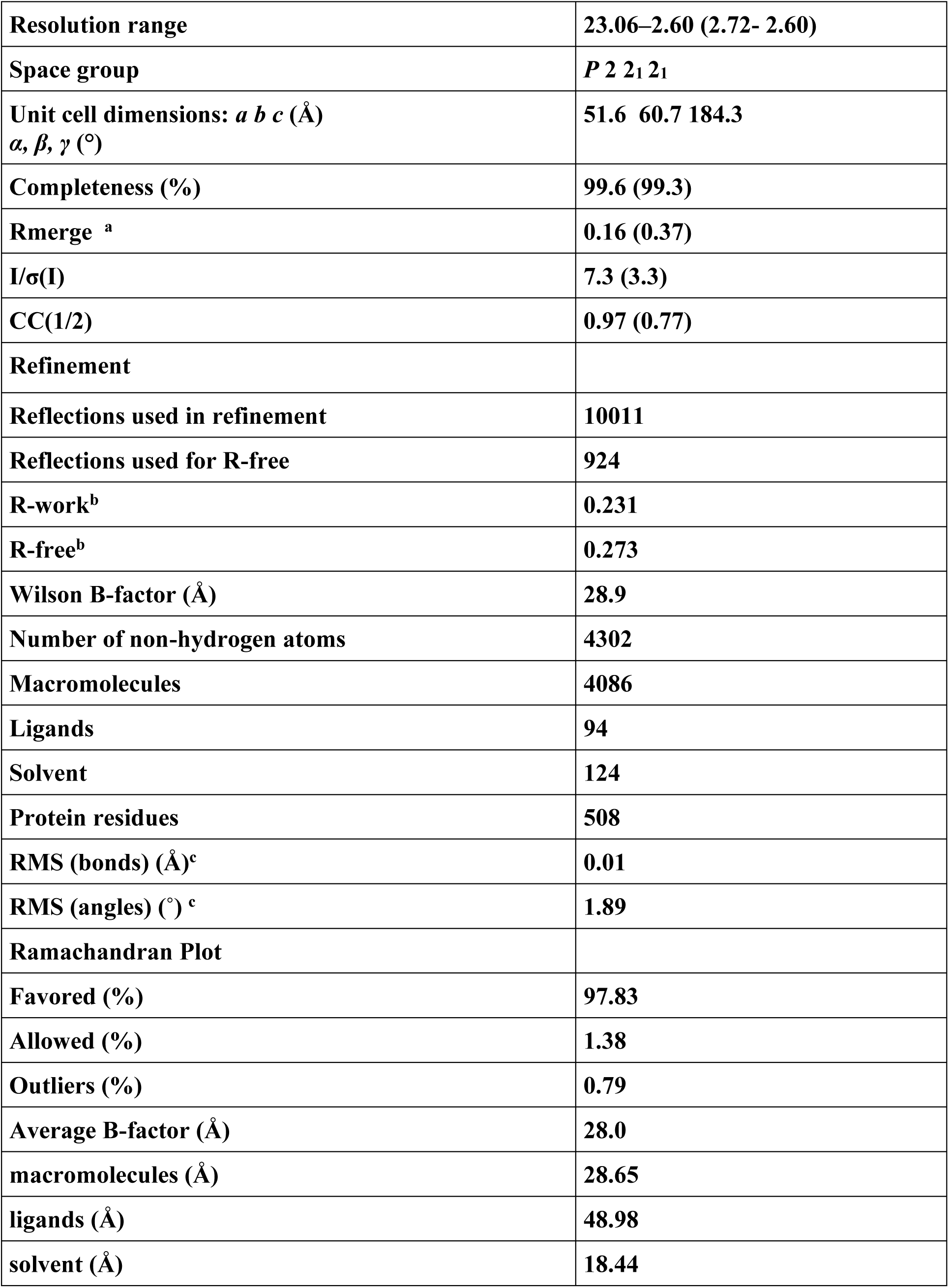
Data collection, processing and refinement statistics of DENV 3 NS5 MTase complexed with soaked CAPE (PDB:8DKZ). Values in parentheses are for the highest- resolution shell. a Rmerge = Σ| I - 〈I〉|/ΣI. b R = Σ|Fobs| - |Fcalc|/Σ|Fobs|. The Rfree is the R calculated on the 5% reflections excluded for refinement. c RMS is root mean square.

The dynamic stability and conformational changes in the Apo proteins and protein-ligands complexes were studied by analyzing their RMSD values during the MD simulation. The average RMSD values for nsP1 Apo was 0.35 nm, and for nsP1 complexes nsP1-SAM, nsP1- HC, nsP1-GTP, and nsP1-CAPE were 0.53 nm, 0.42 nm, 0.46 nm and, 0.38 nm respectively (supplementary figure 3A). Similarly, the Average RMSD for NS5 MTase Apo was 0.23 nm and the NS5 MTase complexes NS5 MTase-GTP, NS5 MTase-HC, and NS5 MTase-CAPE were 0.18 nm, 0.25 nm, and 0.20 nm respectively (supplementary figure 3B). These observations depict that the binding of molecules to nsP1 and NS5 proteins resulted in the formation of a stable complex.

### Binding of compounds to DENV 3 NS5 MTase and CHIKV nsP1

The interactions of NS5 MTase and nsP1 with SAM, GTP, HC, and CAPE at various concentrations were examined using TFS. The intrinsic fluorescence of the native NS5 MTase and nsP1 proteins was measured utilizing a spectrofluorometer. Both proteins showed intrinsic fluorescence quenching with increasing concentrations of all compounds (Figure 3). In TFS, a red shift, indicats to a shift to longer wavelengths, suggests increased polarity or decreased hydrophobicity around the tryptophan residues. Conversely, a blue shift, reflecting a move to shorter wavelengths, signifies reduced polarity or increased hydrophobicity in the local environment (72). A dose-dependent red shift was observed in both proteins upon interaction with HC and CAPE (supplementary figure 4). For nsP1 interactions with SAM, GTP, HC, and CAPE, the K_D_ values were determined to be 10.88 ± 5.15 µM, 169.1 ± 21.31 µM, 6.52 ± 0.55 µM, and 30.57 ± 5.33 µM, respectively (Figure 3 A-D). Similarly, for NS5 MTase interactions with SAM, GTP, HC, and CAPE, the K_D_ values were determined to be 6.67 ± 2.57 µM, 3.03 ± 1.0 µM, 13.35 ± 2.69 µM, and 37.93 ± 6.32 µM, respectively (Figure 3 E-H).

**Figure 3:**
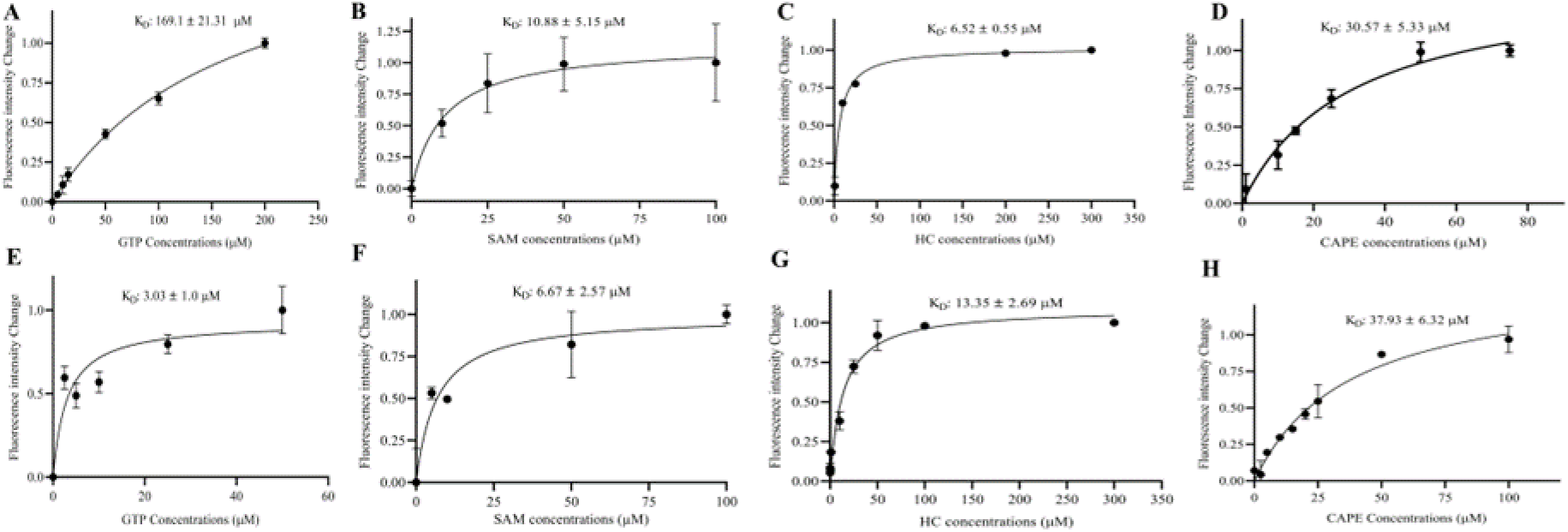
Determination of protein interactions with ligand/substrate by TFS. nsP1 - (A) GTP (B) SAM (C) HC (D) CAPE. NS5 MTase - (E) GTP (F) SAM (G) HC (H) CAPE. Data from three independent experiments were collected and analyzed using nonlinear regression with the ’One Site-Specific Binding’ model.

### Inhibition of Virus-specific MTase by HC and CAPE

To further validate the inhibition of viral MTase activity by HC and CAPE, recombinantly expressed CHIKV nsP1 and DENV NS5 MTase were utilized. Single bands of ∼ 56 kDa and ∼ 32 kDa were observed on 12% SDS-PAGE for purified nsP1 and NS5 MTase, respectively (Figure 4 A,D). The enzymatic activity of nsP1, as assessed through CE-based methods as mentioned in the previously published protocol (14,55). A dose-dependent decrease in MTase activity was observed when treated with HC and CAPE, confirming the significant inhibitory activity of both compounds against CHIKV nsP1 (Figure 4 B,C). Similarly, to validate the inhibition of NS5 MTase activity by HC and CAPE, the CE-based enzymatic activity assay of purified DENV 3 NS5 MTase was performed (14,55). The MTase activity was reduced in response to increasing compound concentrations, confirming the inhibitory effects of HC and CAPE against DENV 3 NS5 MTase (14,55) (Figure 4 E,F).

**Figure 4:**
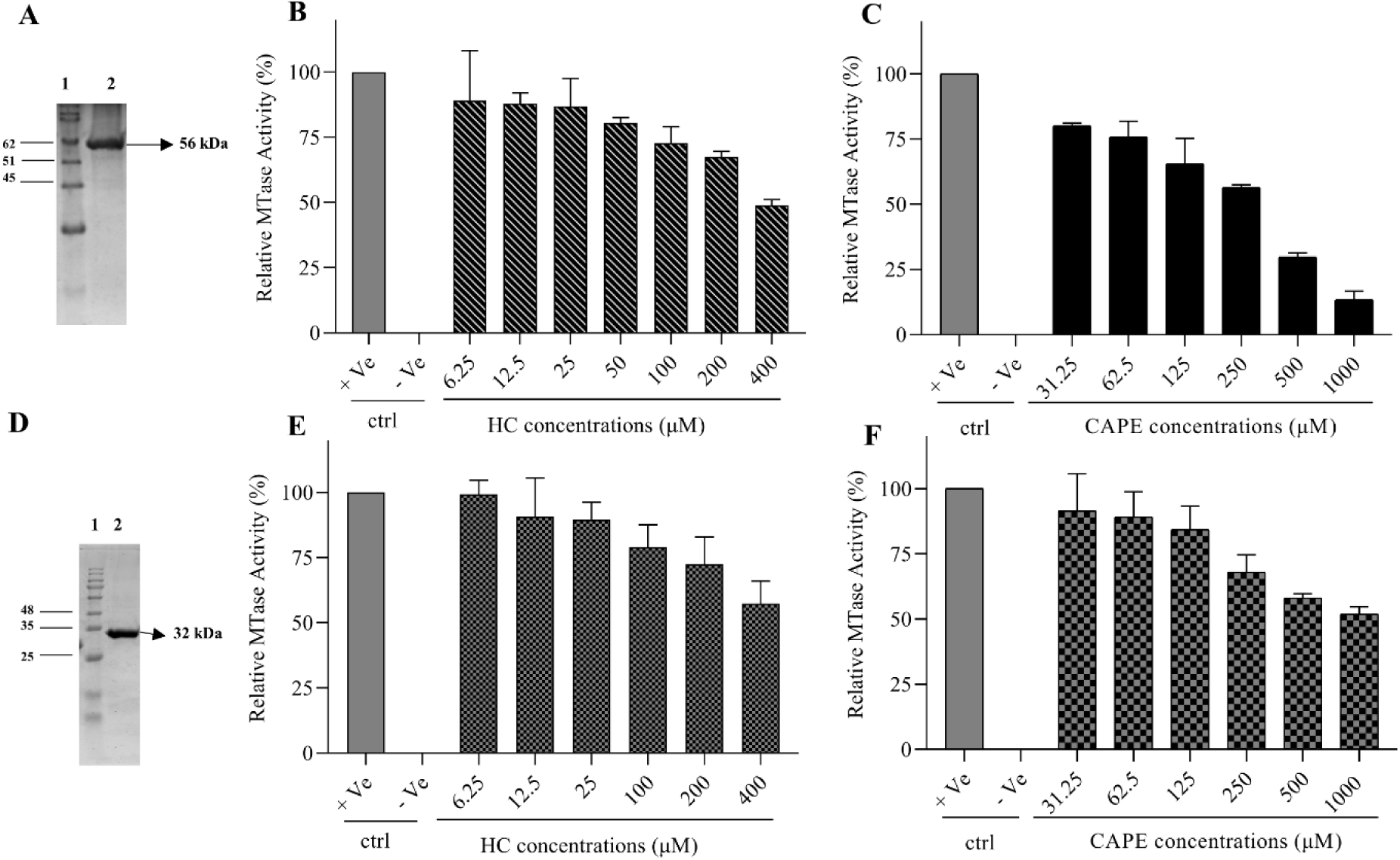
MTase inhibition activity assay. SDS/PAGE analysis of purified recombinant CHIKV nsP1 (A) and DENV 3 NS5 MTase (D). CE-based nsP1 MTase activity inhibition assay (B) HC (C) CAPE and DENV 3 NS5 MTase (E) HC (F) CAPE. Error bars indicate the standard error derived from three independent experiments.

### Structural insights into DENV 3 NS5 MTase with CAPE

The structure of DENV 3 NS5 MTase-CAPE complex (PDB:8KDZ) at 2.6 Å was determined to identify the residues involved in binding to the CAPE molecule (Table 3). Although no SAH was present during crystallization, it was found in both chains, indicating that it has originated from *E. coli* during protein expression (PDB:8KDZ). The main chain conformation in the DENV 3 NS5 MTase-CAPE complex is nearly identical to the DENV 3 NS5-SAH-GTP structure (PDB:4V0R) with RMSD 0.367 Å over 225 Cα atoms (Figure 5C). Two CAPE (302,303) molecules could be modelled in one chain of DENV 3 NS5 structure that matched the observed electron density with a real space correlation coefficient (RSCC) of 0.71 and 0.81, respectively (Figure 5B). The detailed analysis determined by PyMOL and LigPlot^+^, as outlined in Tables 4 and 5, reveals the formation of hydrogen bonds and specific hydrophobic interactions, which are pivotal for the GTP binding site. Arg 211 has an important role in interacting with the phosphate group of GTP (67) (Figure 5E). In CAPE (302), O4 of the 3,4- dihydroxy phenyl group forms a hydrogen bond with NH2 of Arg 211 at 3.5 Å. 3,4-dihydroxy phenyl group of both CAPE form π- π T-shaped interaction with one another. CAPE (302) forms π-σ interaction with Thr 214. Also, its phenyl ring forms π-alkyl interactions with Leu 17, Pro 152. CAPE (303) 3,4-dihydroxy phenyl group acts as π donor to nitrogen of Ser 150 (Figure 5D). Furthermore, CAPE (302) hydrophobically interacts with Asn18, Leu17. These residues are essential in binding the initial nucleotides from octameric RNA (specifically guanosine triphosphate adenosine (G3A) to DENV 3 NS5 MTase (73) (Figure 5F) (Table 6). The CAPE (302-303) molecules also hydrophobically interact with residues Ser150, Thr214, Lys180, and Gly148. Among these residues, Ser150, Thr214, and Lys180 are crucial for hydrogen bonding, while Gly148 is important for the hydrophobic interaction of the adenosine (A)-guanosine (G) dinucleotide from cap 0 RNA to DENV 3 NS5 MTase (22) (Figure 5G, H) (Table 7). Therefore, the presence of CAPE at this position could potentially impede the binding of viral RNA to the DENV 3 NS5 MTase.

**Figure 5:**
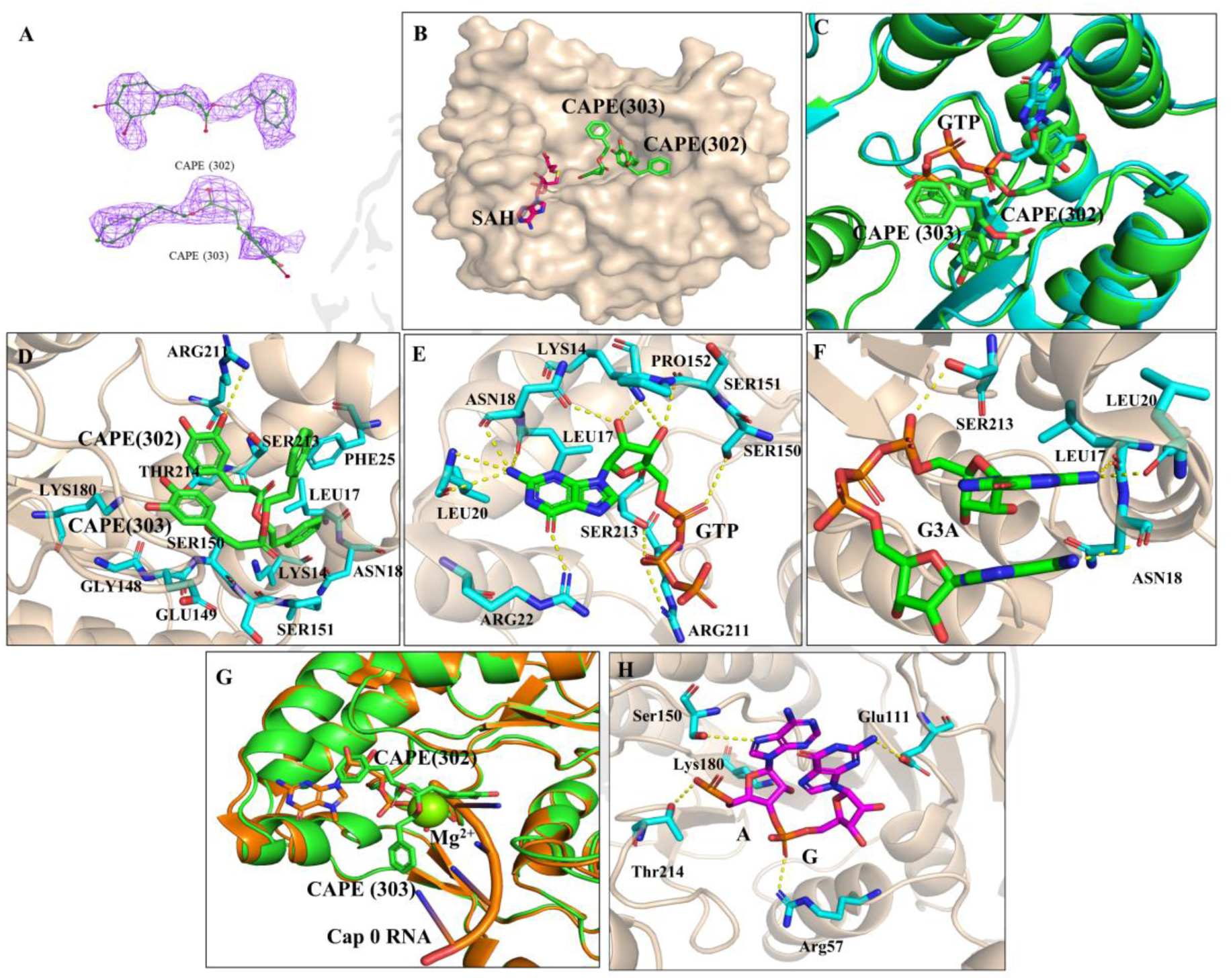
Crystal Structure of DENV 3 NS5 MTase in Complex with SAH and CAPE (PDB:8DKZ). A) Polder omit Map CAPE (302) and CAPE (303) at 3 and 2.5 σ. B) Surface representation of DENV 3 NS5 MTase (wheat) with CAPE (green carbon atoms) and SAH (hot pink carbon atoms) depicted as sticks. C) Structural superposition of a (PDB:8DKZ) green and (PDB:4V0R) cyan. D) Structure of DENV 3 NS5-MTase bound to SAH and CAPE, with hydrogen bond interacting residues represented as yellow dotted lines. E) Structure of the DENV 3 NS5-MTase domain (PDB: 4V0R) bound to GTP, with hydrogen bond interacting residues shown as yellow dotted lines. F) Structure of the DENV 3 NS5-MTase domain (PDB: 2XBM) bound to G3A, with hydrogen bond interacting residues shown as yellow dotted lines. G) Structural superposition of a (PDB:8DKZ) green and (PDB:5DTO) orange. Here, Mg2+ is displayed as a green sphere H) Structure of DENV 3 NS5-MTase bound to A and G from cap 0 RNA, with hydrogen bond interacting residues represented as yellow dotted lines.

**Table 4.**
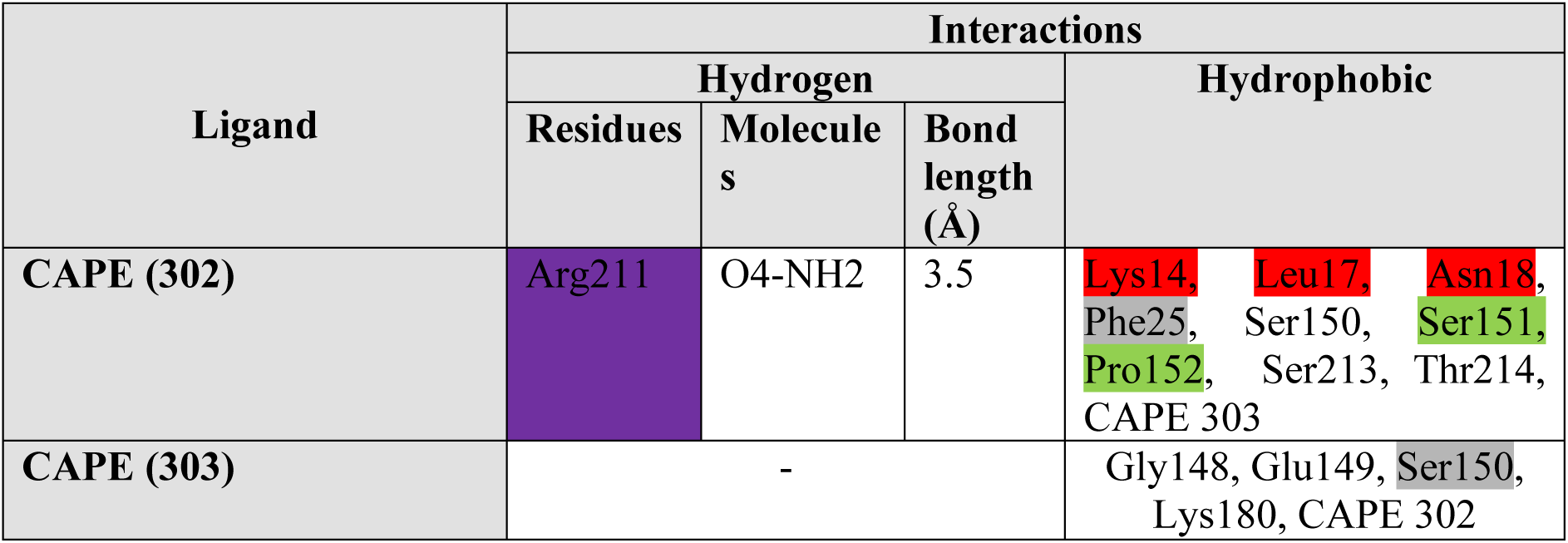
Detailed molecular interactions between the DENV 3 NS5 MTase and CAPE (PDB:8DKZ).

**Table 5.**
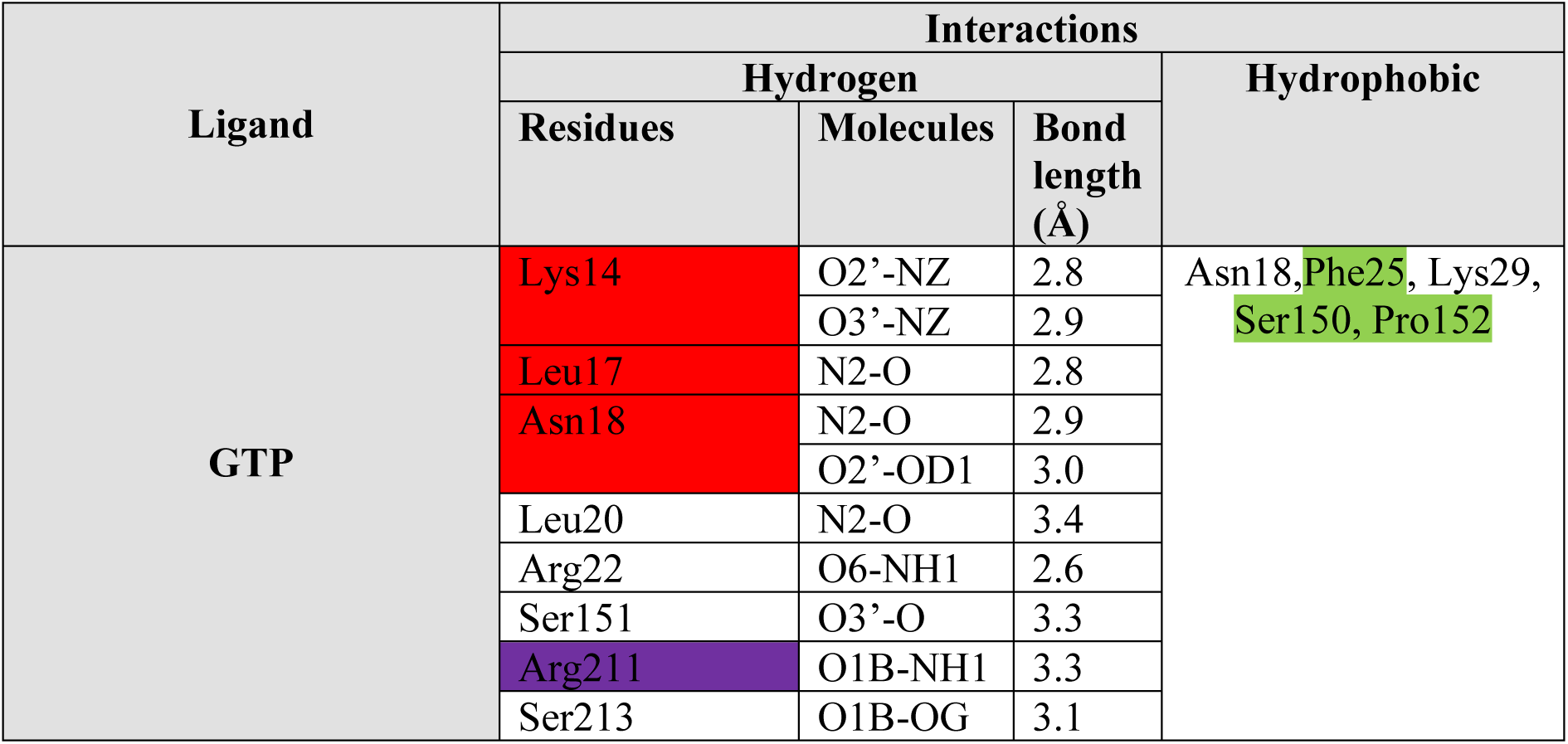
Detailed molecular interactions between the DENV 3 NS5 MTase domain and GTP (PDB:4V0R).

**Table 6.** Detailed molecular interactions between the DENV 3 NS5 MTase domain and G3A (PDB:2XBM).

| Ligand | Interactions |  |  |  |
| --- | --- | --- | --- | --- |
|  | Hydrogen |  |  | Hydrophobic |
|  | Residues | Molecules | Bond length (Å) |  |
| G3A | Asn 18 | O-N11 | 3.26 | Pro 152, Ser 213, Ser 151 |
|  | Leu 20 | O-N32 | 2.81 |  |
|  | Leu 17 | O-N32 | 3.02 |  |

**Table 7.** Detailed molecular interactions between the DENV 3 NS5 MTase domain and A- G (from cap 0 RNA) (PDB:5DTO).

| Ligand | Interactions |  |  |  |
| --- | --- | --- | --- | --- |
|  | Hydrogen |  |  | Hydrophobic |
|  | Residues | Molecules | Bond length (Å) |  |
| A-G | Ser150 | OG-OP1 | 2.88 | Leu 182, Arg84, Lys61, Gly148, Ser56, Glu 216 |
|  | Thr214 | NH2-OP1 | 2.66 |  |
|  | Lys180 | NZ-O2 | 2.35 |  |
|  | Glu111 | OE2-N2 | 2.58 |  |
|  | Arg57 | OG1-OP2 | 3.25 |  |

These color-coded residues represent various interactions in a molecular context. Violet represents hydrogen bonds, green indicates hydrophobic interactions, and red signifies CAPE's hydrophobic residues that make hydrogen bonds with GTP, G3A, and A-G.

### Polyamine-depletion and antiviral activities of HC and CAPE

The MTT colorimetric assay was performed for testing the cytotoxicity of HC and CAPE. The compound concentrations with cell above 90% viability were used subsequently (supplementary figure 5). TLC experiments were performed to determine the effect of HC and CAPE on polyamine levels in mammalian cells treated with these compounds. Vero cells were treated with HC, CAPE, and DFMO, and TLC results confirmed a reduction in the levels of all three polyamines. Put, spd, and spm are positive control markers (Figure 6B). After treatment of Vero cells with CAPE, HC, and DFMO, overall residual polyamine levels are 28.33%, 29.67%, and 46 %, respectively, compared to cell control. Overall, CAPE and HC showed higher polyamine depletion at lower concentrations as compared to the positive control (DFMO) and the cell control (Figure 6B). For this study, the DENV serotype was isolated from a clinical sample, and the DENV 3 serotype was confirmed by PCR using virus and serotype- specific primer pairs and sequencing. The basic local alignment search tool (BLAST) tool revealed 99.78% identity of isolated DENV serotype to DENV 3 isolate NU1883 polyprotein (POLY) gene, partial cds (coding sequences). Similarly, CHIKV (Accession No. KY057363.1.) was propagated in Vero cells and used in further study. Antiviral activities of HC and CAPE at various concentrations were determined against CHIKV and DENV in plaque reduction assays performed in Vero cells using each compound’s non-cytotoxic concentrations. By comparing the viral titer of untreated CHIKV-infected cells with HC- and CAPE-treated CHIKV-infected cells, a dose-dependent decrease in CHIKV titer was observed for HC and CAPE, with IC_50_ values of approximately 13.44 ± 3.21 µM and 0.57 ± 0.03 µM, respectively (Figure 6 C, D supplementary figure 6 A, B). Likewise, HC and CAPE treatment to Vero cells has shown a dose-dependent viral titer decrease in the DENV-infected cells was observed for HC and CAPE, with IC_50_ values of approximately 7.24 ± 2.51 µM and 1.01 ± 0.14 µM, respectively (Figure 6 F, G supplementary figure 6 C, D). DFMO, a known ODC inhibitor, was used as a control and showed less antiviral efficacy against CHIKV at much higher concentrations of 1000 µM than HC and CAPE.

**Figure 6:**
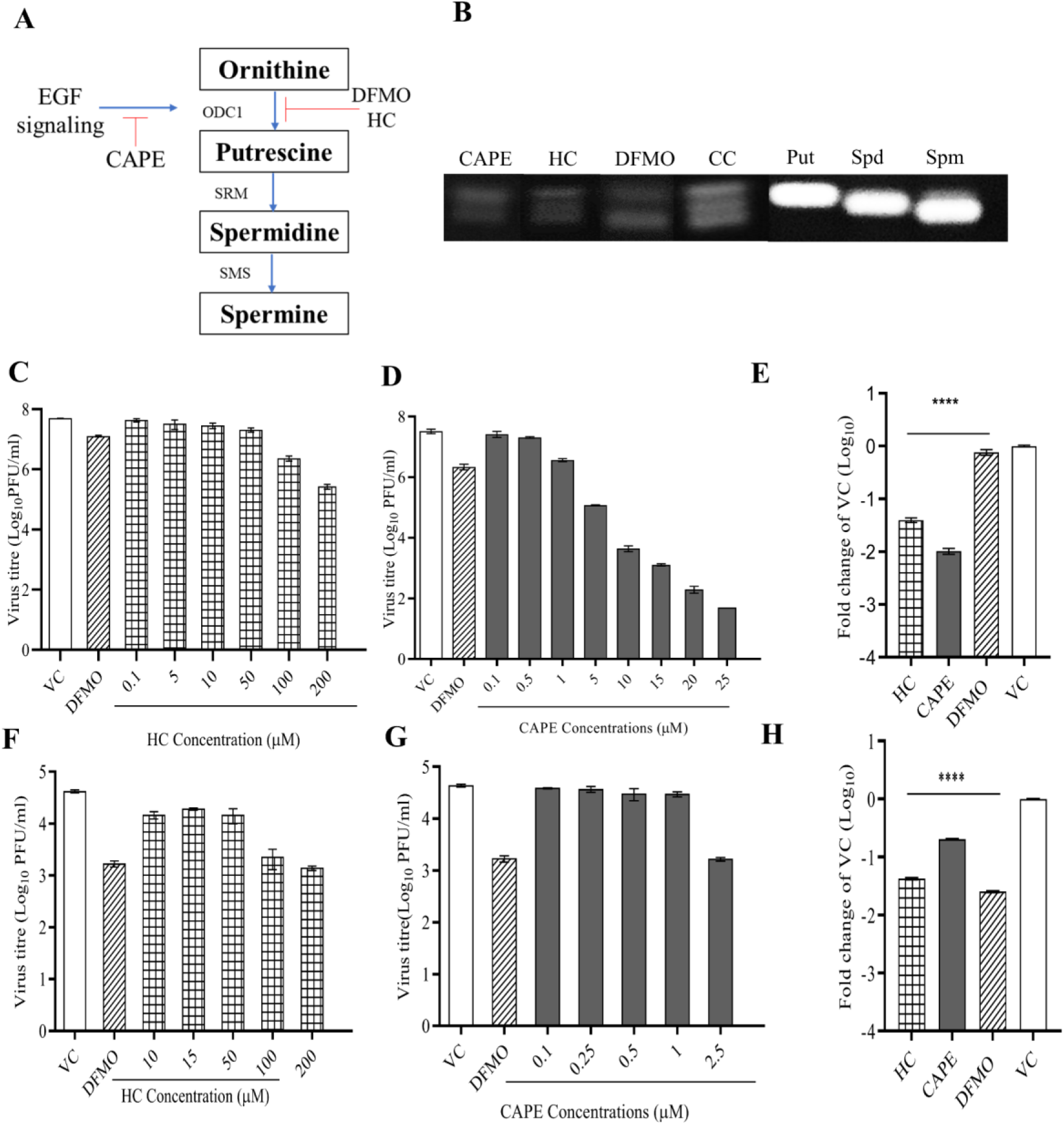
Evaluation of polyamine depletion and antiviral activity using virus titer reduction profiling of HC and CAPE. (A) Mechanistic pathway illustrating the inhibitory effects of CAPE and HC on polyamine pathway and relevant enzymes and inhibitor. (B)the chromatographic analysis of polyamine levels in Vero cells after 36 h treatment with (from left) CAPE (25 µM), HC (200 µM), DFMO(1000 µM), and cell control (CC), 0.1 μM putrescine (Put), spermine (Spm), and spermidine (Spd) as a positive control marker. Treatment with both compounds leads to decreased polyamine levels for both compounds compared to DFMO. Vero cells were treated with HC and CAPE for 12 h and subsequently infected with CHIKV for 2 h. After this, the cells were incubated for an additional 24 h. Following this, the supernatants were collected for plaque assay. A similar compound treatment protocol was followed for DENV infection: Vero cells were pre-treated with the compounds for 12 h, then infected with the virus for 2 h, followed by a 24 h treatment period. After this, the compounds were removed, and the cells were maintained in 2% DMEM and incubated for an additional 4 days. Supernatants were then collected and subjected to plaque assay. DFMO at a concentration of 1000 µM served as a positive control, with a virus control (VC) included for comparison. (C, D, F,G) illustrate the inhibitory effects of various concentrations of HC and CAPE on CHIKV and DENV-infected cells, as assessed by plaque assay. E) RT-PCR for CHIKV with HC 200 µM, CAPE 25 µM, DFMO 1000 µM concentration H) RT-PCR for DENV with HC 200 uM, CAPE 2.5 uM and DFMO 1000 µM. Values are the means, and error bars represent the standard deviation from three independent experiments. Statistical analysis was performed using one-way ANOVA with Dunnett’s post-test. ****P <0.0001.

Here, qRT-PCR was used to validate the antiviral effect of HC and CAPE by quantifying DENV and CHIKV RNA levels in the infected cells. qRT-PCR showed a significant (p < 0.0001) reduction in the viral RNA levels when treated with HC and CAPE compounds in an antiviral assay for CHIKV (Figure 6E) and DENV (Figure 6H). At the mentioned concentrations, HC and CAPE showed a 23- and 4-fold reduction in DENV, respectively, compared to the virus control (VC). Similarly, HC and CAPE reduced CHIKV by 25 and 97 fold at the same doses, respectively, compared to VC. IFA results further corroborated these results. The results of the IFA analysis showed a reduction in CHIKV (Figure 7A) and DENV (Figure 7B) after treatment with HC and CAPE.

**Figure 7:**
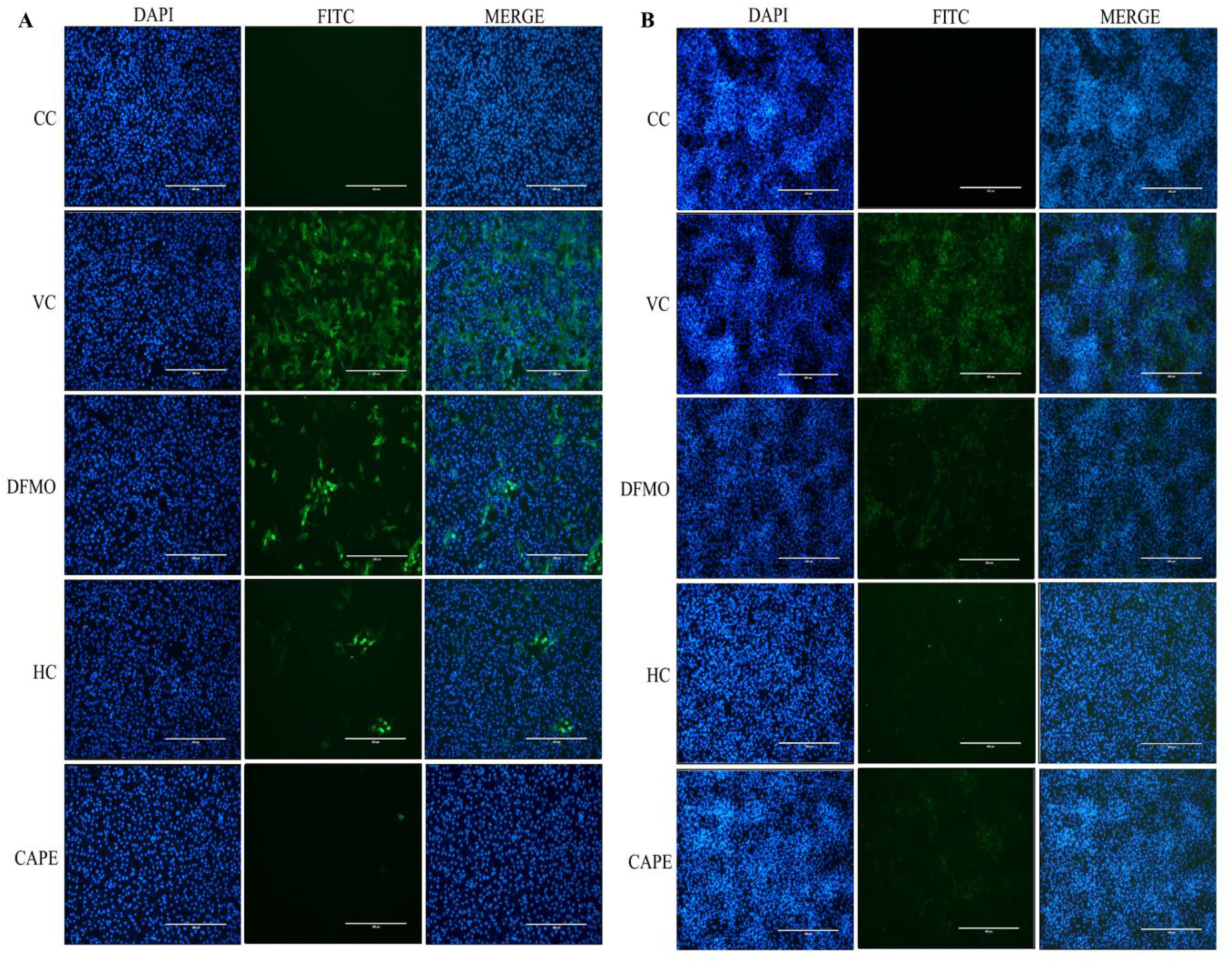
Evaluation of the antiviral effect by immunofluorescence assay (IFA). Immunofluorescent staining of HC (200 µM) and CAPE (25 µM for CHIKV and 2.5 µM for DENV) treated with CHIKV(A) and DENV(B) in Vero cells. Green fluorescence indicates the virus load, and blue fluorescence indicates the nuclear staining with DAPI with a 10 X objective lens. The scale bar is 400 μm.

Further, the effects of the addition of exogenous biogenic polyamines were studied. Unlike DFMO-treated cells, the exogenous addition of all three polyamines at 1 µM were unable to effectively rescue virus titer after CAPE and HC treatment for both CHIKV and DENV (Figure 8).This finding suggests an additional mechanism is involved in the antiviral action of these compounds against both viruses, supporting the initial hypothesis.

**Figure 8:**
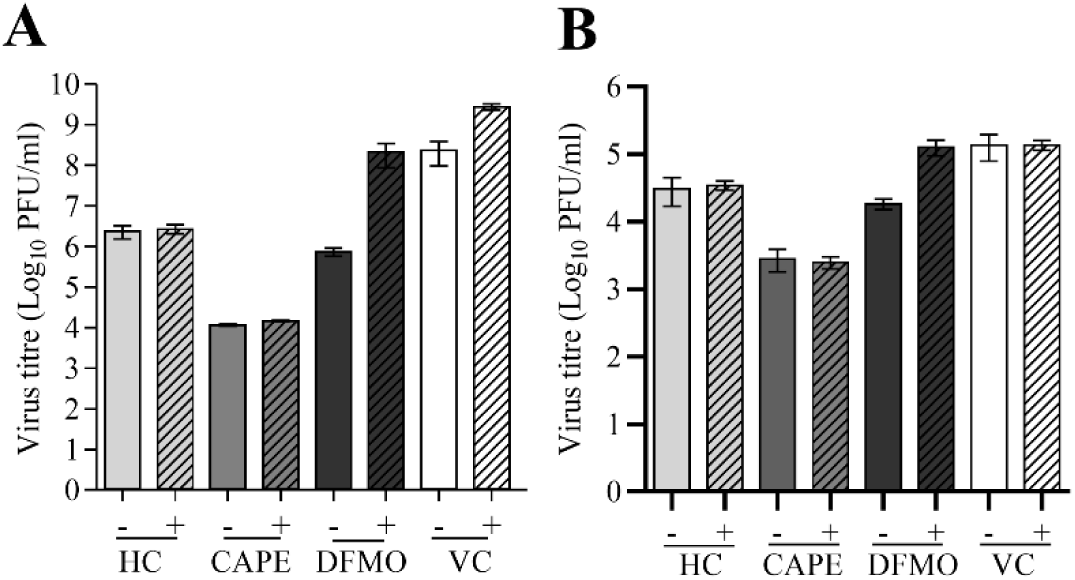
Effect of addition of biogenic polyamines on virus infected and compound treated cells. Vero cells were pre-treated with drugs for 24 h at various concentrations, following this cells were infected with CHIKV (A) and DENV (B) treated with HC (200 µM), CAPE (25 µM for CHIKV and 2.5 µM for DENV), DFMO (1000 µM) as a positive control and VC as untreated control. Minus (-) sign indicates only the compound-treated Vero cells, whereas plus (+)sign indicates that 1µM polyamines (put, spd, spm) were added additionally in the compound-treated cells. Error bars represent the standard error of three independent experiments.

## Discussion

The continuous emergence and re-emergence of viruses like DENV and CHIKV underscores the need for effective public health interventions and antiviral therapeutics to mitigate their spread. Although FDA-approved vaccines for DENV and CHIKV are available, no antiviral therapies are approved against these viral infections (4,7). Developing novel strategies to combat these life-threatening human pathogens, by targeting both viral enzymes and host factors, holds significant promise for accelerating drug development. This dual approach can potentially enhance therapeutic efficacy and circumvent resistance by addressing critical components of viral replication and host-pathogen interactions. Interestingly, SAM and GTP binding sites in nsP1 and the GTP binding site in NS5 MTase are conserved across alphaviruses and orthoflaviviruses infecting humans, insects, and aquatic life (supplementary figure 1 A,B). An *in house* library of NSMT was screened against the CHIKV nsP1 and DENV NS5 MTase. The screening identified HC and CAPE as the top hits with stable interactions to the active sites of both enzymes.

In tryptophan fluorescence spectroscopy, the binding affinity of HC and CAPE with CHIKV nsP1 and DENV NS5 MTase was determined and compared to that of SAM and GTP. SAM and GTP induced fluorescence quenching without spectral shifts, indicating localized changes near tryptophan residues without major conformational alterations (supplementary figure 4) (72). Conversely, both HC and CAPE have demonstrated a dose-dependent red shift with fluorescence quenching, indicating major structural changes upon interaction (Figure 3 and supplementary figure 4). HC and CAPE have shown dose-dependent inhibition in nsP1 and NS5 MTase CE-based MTase assays (Figure 4).

The crystal structure of CAPE in complex with the DENV 3 MTase reveals that two CAPE molecules bind to the GTP and cap 0 RNA binding site (Figure 5). Similarly, structural analysis of NS5 MTase and CAPE (303, 302) complex revealed a similar binding pattern of CAPE molecule with residue Lys14, Leu17, Asn18 (CAPE-303) and Gly148 for (CAPE-302) of NS5 MTase as predicted in docking studies (Table 2,4). Among all hydrophobic interactions of CAPE, residues Leu17, Asn18, Phe25, Ser150, Gly148, Glu149, Ser213, and Lys180 are conserved across the NS5 MTase, as illustrated (supplementary figure 1B) (Table 4). Most of these residues are involved in maintaining structural stability, with Lys180 being part of the Lys61-Asp146-Lys180-Glu216 catalytic tetrad (K-D-K-E). Here, Lys180, acting as the catalytic base, activates the 2’-OH group of the ribose sugar to facilitate nucleophilic attack on the methyl group carbon atom of SAM (22). The CAPE binding site encompasses magnesium (Mg), critical in stabilizing the structural integrity between cap 0 and RNA (Figure 5 G). This is achieved through the hexacoordination of Mg2+ ions, involving three oxygen atoms from phosphate and three from water molecules. Furthermore, Mg2+ ions form hydrogen bonds with residues Ser213 and Ser150 protruding from the protein surface (22). Given its proximity to this site, CAPE has the potential to influence the stability of the cap 0-RNA complex. In OMHV, GMP-enzyme intermediate was identified within the GTP binding site (22). Hence, it is possible that CAPE molecules in the GTP binding site can affect the viral RNA binding and interfere with the capping process. Although this work did not address the possibility of inhibiting viral RNA binding activity, incorrectly capped viral RNA can lead to inefficient translation of viral proteins (20,21). AT-9010 (PDB: 8BCR) and ribavirin triphosphate (PDB: 1R6A) have previously been observed binding to the GTP binding site of the NS5 MTase. Both compounds, which are GTP analogues, exhibit anti 2′-O-MTase activity (Table 8) (74,75). While using nucleoside analogues is a promising antiviral strategy, it can potentially disrupt cellular functions and lead to the rapid development of resistance (76).

**Table 8.**
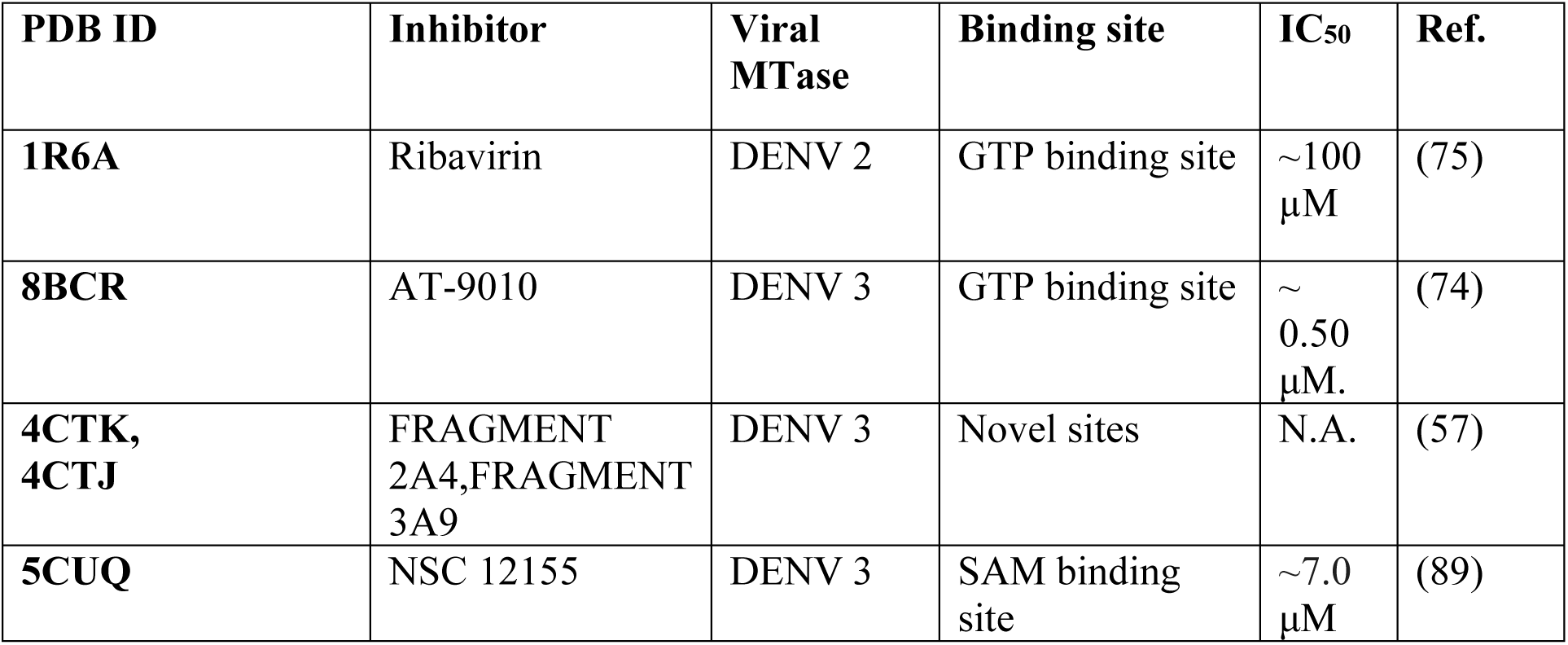
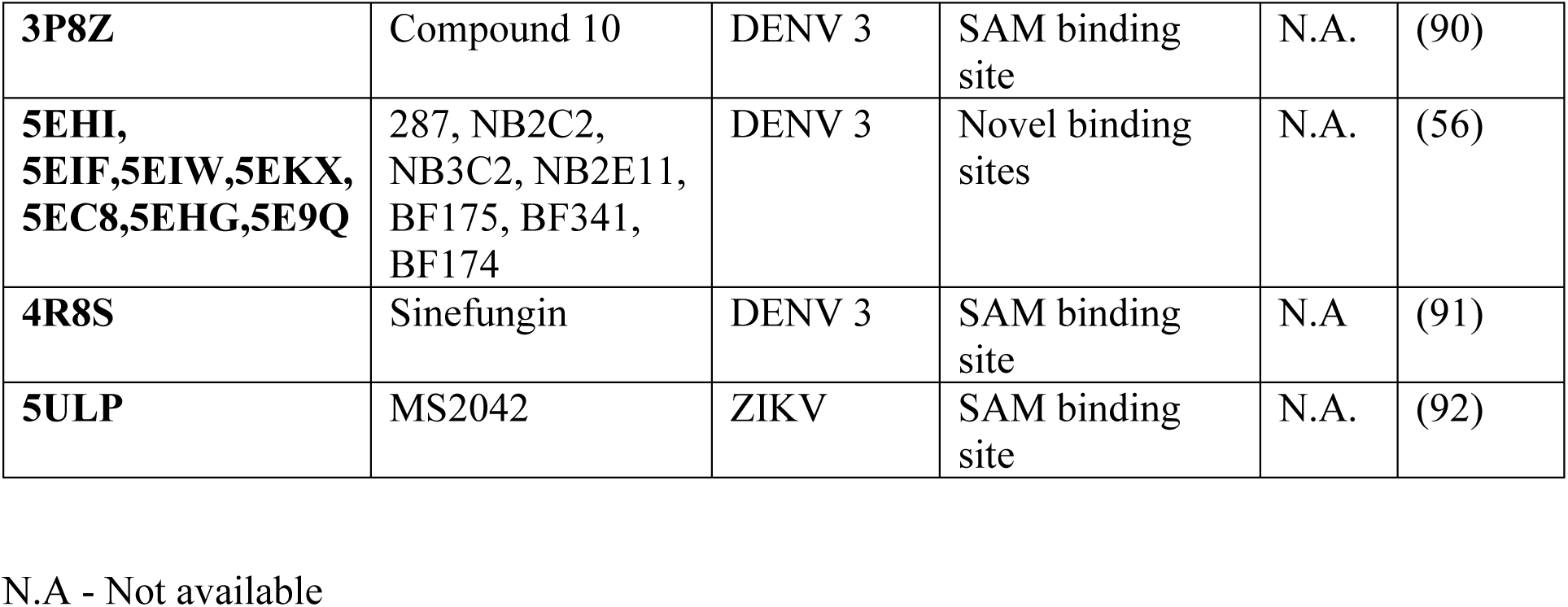
Summary of reported DENV MTase Inhibitors.

Polyamines are ubiquitous molecules present in all cells and play a crucial role in the viral life cycle (77). Polyamine depletion can impair cellular processes crucial for cell survival and homeostasis, such as gene expression, apoptosis, protein synthesis, oxidative stress levels, and cell signaling regulation (78–80). However, DFMO, which inhibits the ODC enzyme of the polyamine biosynthesis pathway, is an FDA-approved drug for African sleeping sickness and high-risk neuroblastoma (81,82). Moreover, DFMO is administered via a short-course regimen (14 days) in African sleeping sickness treatment to mitigate potential adverse effects (83,84). Previous studies showed that DFMO treatment cells depleted biogenic polyamine levels, leading to its antiviral activity against various viruses such as CHIKV, ZIKV, and DENV (25). This study supported a previously published finding that HC and CAPE are polyamine pathway inhibitors (85,86). HC is a naturally occurring flavonoid that is an established ODC inhibitor (86). Flavonoids have significant therapeutic potential as antiviral agents against various classes of viruses (87).

CAPE is a natural ester that has also shown antiviral activity, and it significantly prevented the full induction of ODC by epidermal growth factor (EGF) (85,88). HC and CAPE effectively reduced CHIKV and DENV infections more than DFMO in plaque reduction assays and IFA experiments, validating their superior inhibition in virus-infected cells (Figure 6,7). qRT-PCR quantified viral RNA from CHIKV and DENV-infected cells, reflecting viral replication. HC and CAPE treatment significantly reduced viral load compared to the VC (Figure 6).

This study shows HC and CAPE molecules have shown better polyamine depletion than the DFMO (Figure 6B). An exogenous polyamine addition assay was conducted following the methodology outlined by Mounce et al. (2016), adhering to established procedures. Consistency was observed between the results of virus titer rescue in DFMO treatment upon exogenous polyamine addition. However, unlike DFMO treatment for HC and CAPE, the exogenous polyamine addition assay did not rescue viral titer for CHIKV and DENV (Figure 8 A, B). The inability of external polyamine supplementation to restore viral titers implies that polyamine depletion is not the sole mechanism of action. Therefore, the data supports the hypothesis of an additional inhibitory mechanism, likely involving targeting viral MTases. These novel natural antiviral molecules and the findings of this study will further assist in developing broad-spectrum antiviral strategies against emerging alphaviruses and orthoflaviviruses.

## Conclusion

This research provides compelling first evidence of HC and CAPE’s anti-CHIKV and anti-DENV activities through two mechanisms: a) indirectly by depleting polyamines in mammalian cells and b) directly by targeting viral MTases. The detailed atomic interactions revealed that CAPE binding at the GTP and cap 0 RNA binding sites in DENV MTase may impede viral RNA capping mechanism. HC and CAPE’s novel broad-spectrum antiviral activity makes them promising candidates for developing anti-DENV and anti-CHIKV therapeutics.

## Acknowledgments

The authors thank the Department of biosciences and bioengineering, IIT Roorkee for central facilities and the translational and structural bioinformatics center. The authors thank the Macromolecular Crystallographic Facility (MCU) at IIC, Indian Institute of Technology, Roorkee. ST and PK thank the Department of Biotechnology, Govt of India, for supporting Bioinformatics Center at IIT Roorkee ref number BT/PR40141/BTIS/137/16/2021. ST and PK also would like to thank Department of Biotechnology, Govt of India “National Network Project of Department of Biotechnology, Indian Institute of Technology, Roorkee” Project no. BT/PR40142/BTIS/137/72/2023. The authors would like to acknowledge Mr. Mayur Ghate, Ms. Bharati for technical assistance in TFS analysis. The authors would like to thank the University Grants Commission (UGC), Council of Scientific and Industrial Research (CSIR), Indian Council of Medical Research (ICMR), and Ministry of Human Resource Development (MHRD) for providing financial support. This work was supported by a research grant to ST from ICMR ref no. ISRM/12(46)/2020.

## Ethical approval

DENV-positive clinical sample was collected from HiTech Pathology Laboratory, Roorkee India with the approval of the Institutional Human Ethical Committee (IHEC), Indian Institute of Technology, Roorkee (Ref no. BT/IHEC-IITR/2016/6499).

## Supplemental information

Supplemental information includes supplemental experimental procedures and six Figures.

## Authors contribution

MB, AK, RR, VS, AP, AK, SM, BW, and RK conducted the experiments. MB, AK, RR, VS, RM, DS, PK, and ST planned the experiments and analyzed the data. MB, PK, and ST took the lead in writing the manuscript. All authors provided critical feedback and helped shape the research, analysis, and manuscript.

## Competing interests

The authors declare that they have no competing interests.

## Supplementary Experimental Procedures

### Cell line, Virus isolation, propagation and serotyping

Vero cell line isolated from kidney epithelial cells of an African green monkey was obtained from the National Centre for Cell Science (NCCS), Pune, India. Vero cells were used for the propagation and titration of DENV and CHIKV. The cells were maintained in Dulbecco’s Modified Eagles Medium (DMEM) (Gibco, HiMedia) supplemented with 10% inactivated fetal bovine serum (FBS) (Gibco, HiMedia) along with 100 U/mL penicillin and 100 mg/mL streptomycin (Gibco, HiMedia). The cells were maintained at 37°C with 5% CO_2_ supplementation.

Dengue was isolated from the DENV-suspected patient’s blood samples. Sera was aliquoted and tested for the presence of DENV NS1 antigen by J Mitra ELISA kit according to the manufacturer’s instructions. DENV-positive samples were used for virus isolation. Initially, sera were 1:10 diluted in minimum essential medium (MEM) (Gibco, HiMedia) containing 2 % FBS, 100 U/mL penicillin, and 100 mg/mL streptomycin for virus propagation. Then, Serum samples were added to a confluent monolayer of Vero cells for 90 min with gentle shaking every 15 min in a 37 °C CO_2_ incubator. The inoculum was withdrawn, and the cells were cultured for 5-6 days in fresh MEM with 2% FBS and antibiotics until the appearance of cytopathic effect (CPE). After 5-6 days, the supernatant was harvested and serially diluted for infection of Vero monolayer cells for 90 min and overlaid with MEM containing 1 % agarose for the plaque purification procedure. Single plaques were picked and resuspended in MEM and were further propagated in Vero cells. Then, DENV NS1 ELISA-positive supernatants showing CPE was collected and used for further study.

To confirm the DENV and its serotype, RNA from the supernatant of cell culture infected with the virus was isolated using the Trizol (Sigma) method described by the manufacturer. The RNA was then reverse-transcribed into complementary DNA (cDNA) using the PrimeScript cDNA Synthesis Kit (Takara). Synthesized cDNA was utilized as a template for polymerase chain reaction (PCR) (63) and the amplification product was analyzed by gel electrophoresis in 1 % agarose gels stained with ethidium bromide. The sample was sent for DNA sequencing, and the serotype of DENV was confirmed by matching the obtained sequence against the nucleotide database.

CHIKV (Accession No. KY057363.1.) was propagated and titrated in a Vero cell line using the protocol reported by Singh et al., 2018 and then stored at −80 °C for further experiments (61).

### Cell Cytotoxicity Assay

Different concentrations of HC and CAPE were evaluated for cytotoxicity on Vero cells using 3-(4,5-dimethyl-thiazol-2-yl)-2,5-diphenyltetrazolium bromide (MTT) (Himedia) assay. Before treatment, Vero cells were seeded in a 96-well plate, and at 90% confluency, the media were removed, and different dilutions of compounds were added into each well for 12 h. After incubation, the inoculum was removed, and fresh maintenance media were added for 2 h. Further fresh compound dilutions were added to each well and kept for 24 h. Subsequently, 20 μL/well of MTT (5 mg/mL) was added and incubated for 4 h at 37 °C in 5% CO_2_. Upon incubation, 110 μL/well of DMSO was added to dissolve formazan crystals. Plates were read at a wavelength of 570 nm using cytation 3 multi-mode plate reader (BioTek Instruments, Inc.). The average absorbance of 0.1 % DMSO-treated cells was used as cell control. In the second set of experiments, after completion of 24 h post-treatment, fresh maintenance media were added and kept for 4 days. The cell viability of the treated well was compared with the cell control, and the concentration that showed >90% viability after compound treatment was considered non-toxic.

### Plaque assay

Vero cells were seeded at 1.0 x 10^5^ cells/well in a 24-well plate in complete media before infection. The supernatant was 10-fold serially diluted in maintenance media and was inoculated on ∼80-90% confluent cells. Plates were incubated for 2 h with gentle shaking every 15 min for virus adsorption at 37 °C with a 5% CO_2_ incubator. After adsorption, overlay media and maintenance media were added in 1:1 dilution and plates were further incubated for 2 days and 3 days for CHIKV and DENV, respectively at 37 °C in 5% CO_2_ incubator. Following that, cells were stained with 1% crystal violet to count the number of CHIKV plaques and immunostaining for DENV-infected cells, as described below.

Immunostaining was performed at room temperature. After incubation, overlay media was removed from the plates, and the cell monolayer was fixed with 3.7% formaldehyde solution for 30 min. Cells were washed 3 times with PBST (0.02% Tween-20 in Phosphate buffer saline). After that, cells were permeabilized with 0.2% triton X-100 in PBS for 7 min. Cells were washed three times with PBST and incubated with 3% Skim milk for 30 min. Then, cells were washed three times with PBST and incubated with 1:500 diluted orthoflavivirus group antibody (Genetex) for 2 h. Cells were washed three times with PBST and incubated with secondary antibody (Goat anti-mouse IgG HRP, 1:1500 dilution) for 1 h. After washing 2 times with PBST and three times with PBS, cells were stained with True Blue Peroxidase substrate (KPL, Sera Care, MA, USA) and incubated in the dark for 30 min to develop blue color staining of virus-infected cells, and foci were counted (62).

### Multiple Sequence Alignment (MSA)

The amino acid sequence of nsP1 protein of *alphaviruses* was compared with CHIKV nsP1 as a reference point using Clustal Omega (40) to check if the key residues involved in the capping of viral RNA were conserved across different viruses. The MSA was performed for Venezuelan equine encephalitis virus (VEEV Gene Bank Id: AAU89533.1), (CHIKV Gene Bank Id: QOW97289.1), Ross River virus (RRV Gene Bank Id: QTC33397.1), Sindbis virus (SINV Gene Bank Id: AWT57896.1), Aura Virus (AURA Gene Bank Id: AWQ38330.1), Middelburg virus (MIDV Gene Bank Id: QOY44469.1), Barmah Forest virus (BFV Gene Bank Id: QVM79755.1), Madariaga virus (MADV Gene Bank Id: ABL84688), Salmonid alphavirus (SAV Gene Bank Id: KC122922.1), Eilat virus (EILV Gene Bank Id: NC_018615) and Mayaro virus (MAYV Gene Bank Id: AZM66145) from the alphaviruses. The sequence alignment profile of the selected nsP1 sequences was performed via Clustal Omega tool and analyzed by a graphical coloured depiction using ESPript 3.0 (40).

Similarly, MSA was performed for the MTase domain of NS5 from dengue 3 Virus (DENV 3 PDB Id:4CTJ), ZIKV PDB Id:5WZ2, West Nile Virus (WNV PDB Id:2OY0), Yellow Fever Virus (YFV PDB Id:3EVA), Palm Creek virus (PCV Gene Bank Id: NC_033694.1), Wenzhou shark flavivirus (WSF Gene Bank Id: AVM87250.1) and Japanese Encephalitis Virus (JEV Gene Bank Id:4K6M_1) from the orthoflaviviruses and the analysis was carried out in the same way as for DENV.

## Abbreviations

HC: Herbacetin
CAPE: Caffeic acid phenethyl ester
CHIKV: Chikungunya virus
DENV: Dengue virus
MTase: Methyltransferase
NS5: Nonstructural protein 5
nsP1: Nonstructural protein 1
DAPI: 4’,6-diamidino-2-phenylindole
qRT-PCR: Quantitative reverse transcription polymerase chain reaction
PFU: Plaque-forming units
GTP: Guanosine triphosphate
SAH: S-adenosyl homocysteine
SAM: S-adenosyl methionine

---

**Supplementary Figure 1:**
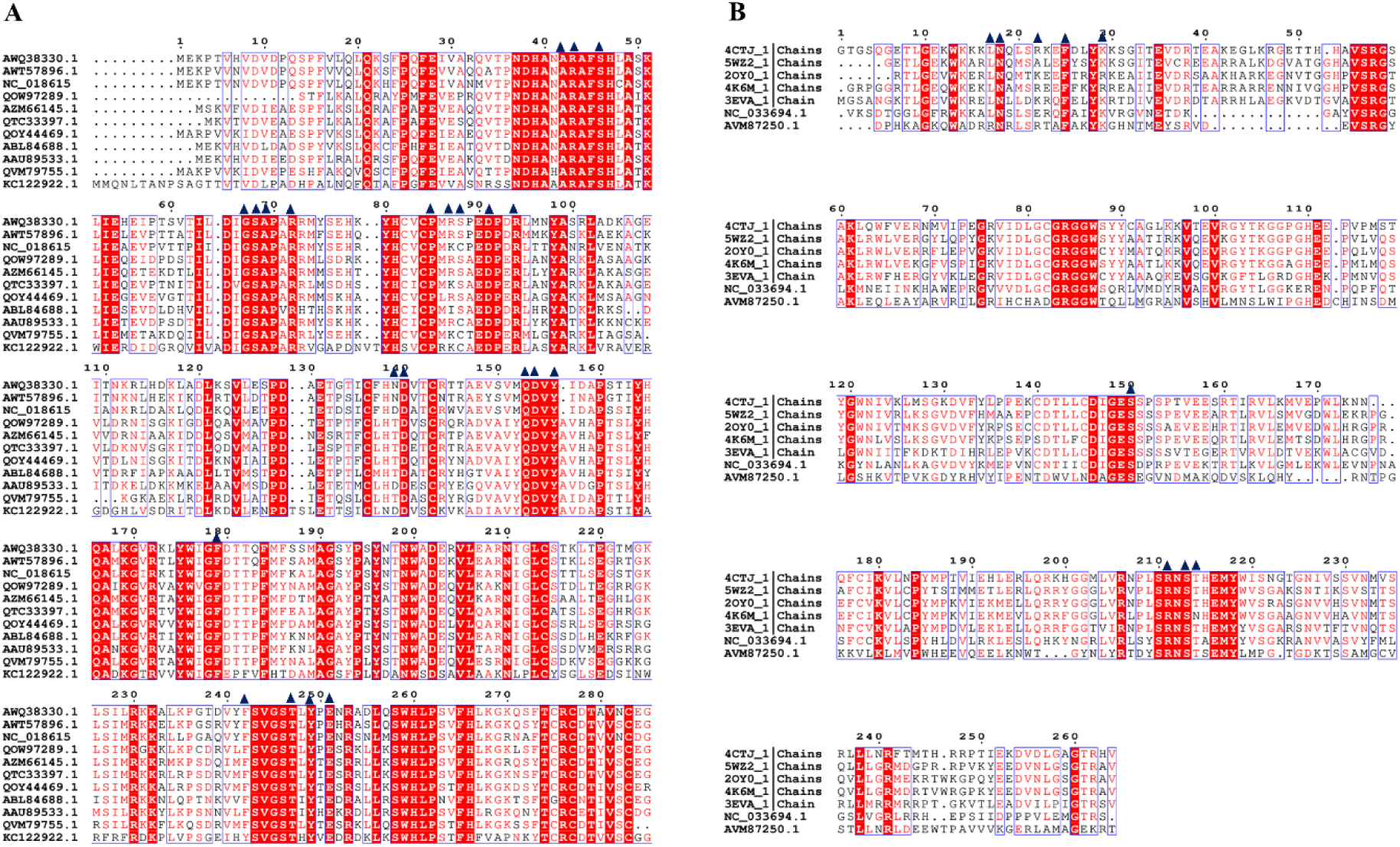
Multiple Sequence alignment of MTase domains of different alphaviruses and orthoflaviviruses. A) Sequence alignment of nsP1 MTase domain (residues 1 to 285) from CHIKV with the MTase domains of other alphaviruses. VEEV Gene Bank Id: AAU89533.1, CHIKV Gene Bank Id: QOW97289.1, RRV Gene Bank Id: QTC33397.1, SINV Gene Bank Id: AWT57896.1, AURA Gene Bank Id: AWQ38330.1, MIDV Gene Bank Id: QOY44469.1, BFV Gene Bank Id: QVM79755.1, MADV Gene Bank Id: ABL84688, SAV Gene Bank Id: KC122922.1, EILV Gene Bank Id: NC_018615 and MAYV Gene Bank Id: AZM66145. B) Sequence alignment of NS5 MTase domain (residues 1 to 264) from DENV with MTase domains of other orthoflaviviruses. DENV 3 PDB Id:4CTJ, ZIKV PDB Id:5WZ2, WNV PDB Id:2OY0, YFV PDB Id:3EVA, PCV Gene Bank Id: NC_033694.1, WSF Gene Bank Id: AVM87250.1 and JEV PDB Id:4K6M. The active-site MTase residues in both alphaviruses and orthoflaviviruses are highly conserved, suggesting that the structure and function of the MTase domain in both families are conserved. Residues are represented with single letter amino acid code, with identical residues indicated in white font and boxed in red, while similar residues are indicated in red font. Blue triangles correspond to the MTase active site residues as per the literature. Alignments were generated via Clustal Omega and ESPript tools.

**Supplementary Figure 2:**
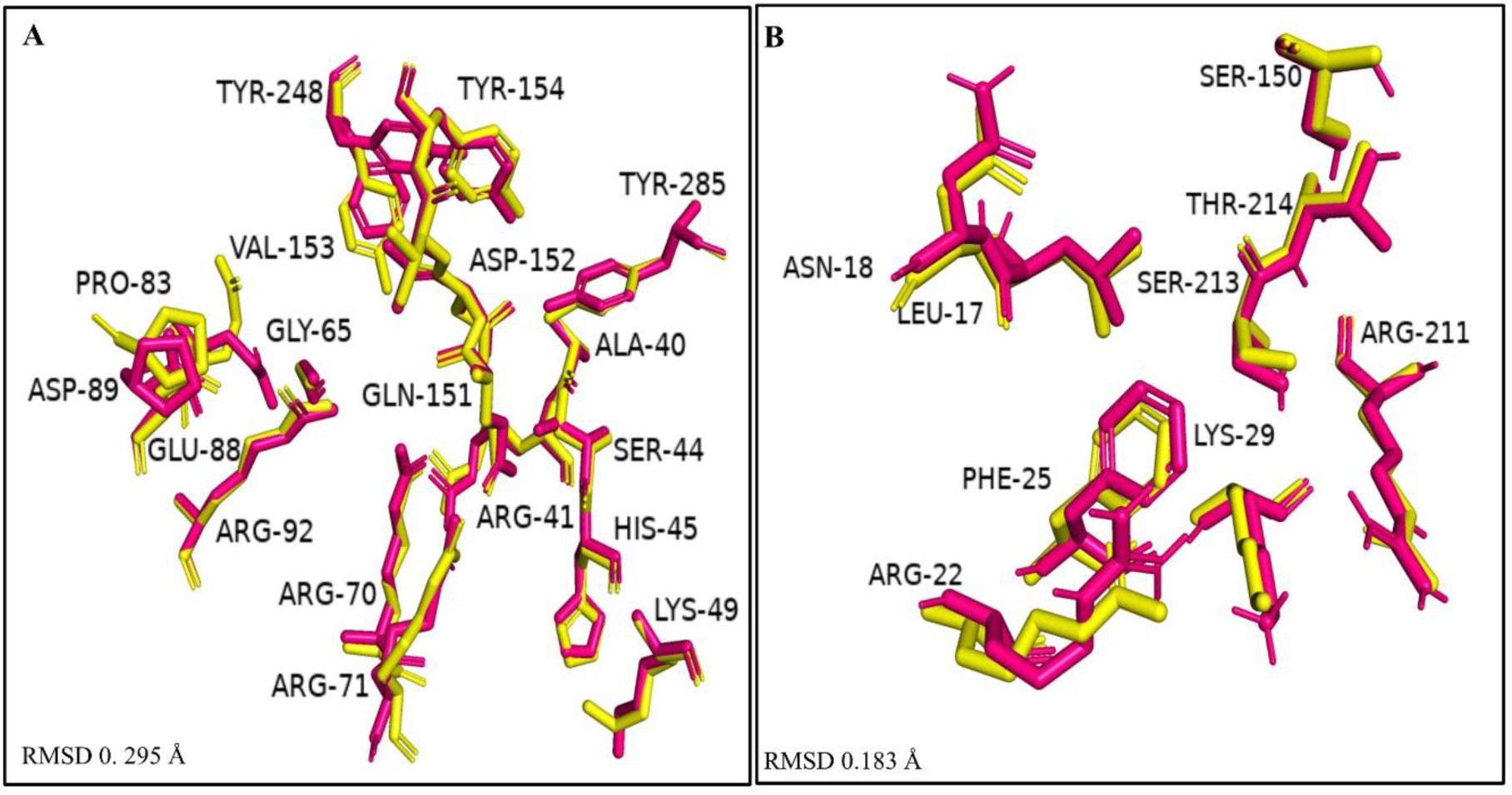
Superposition of active site residues of SWISS generated models with templates. (A-B) Close-up view of overlapped residues of generated models of CHIKV nsP1 (A) and DENV 3 NS5 MTase (B) with RMSD values of 0.295 Å and 0.182 Å, respectively. The generated model is labelled yellow, and the template is labelled pink.

**Supplementary Figure 3:**
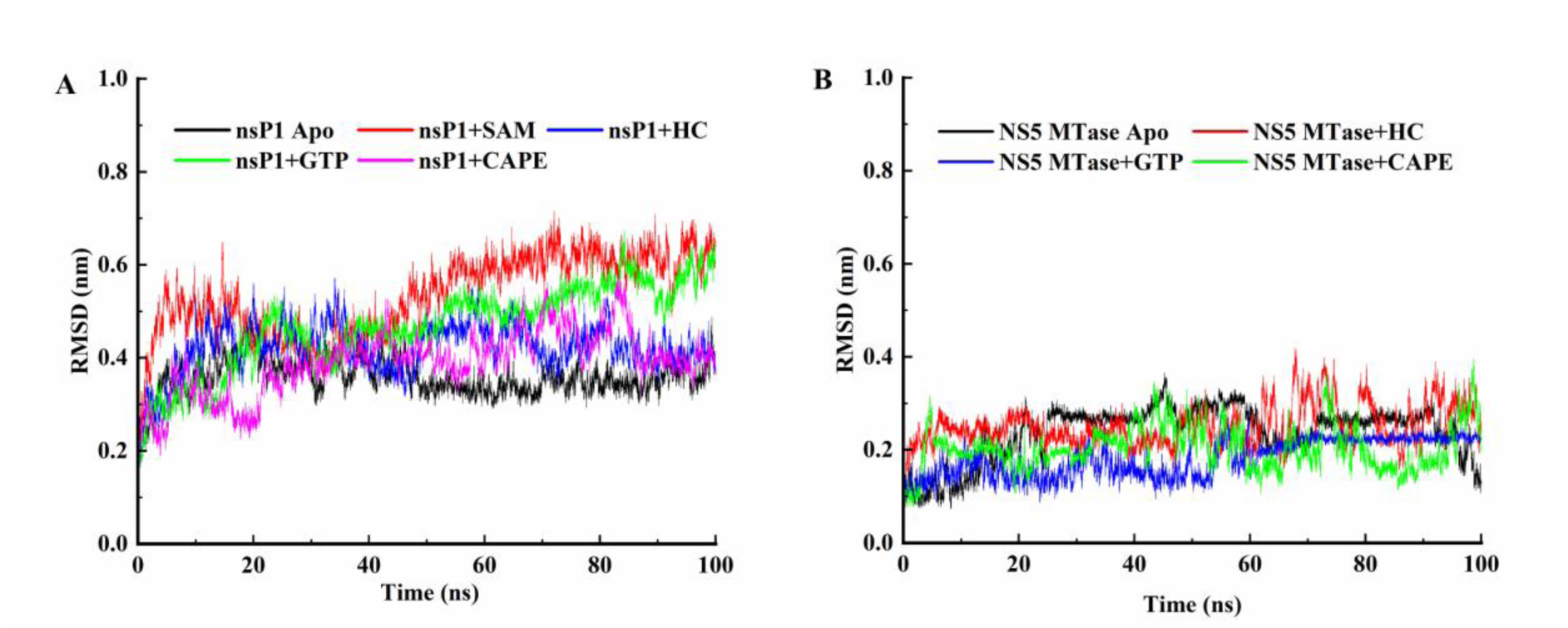
Root Mean Square Deviations (RMSD) graphs of (A) CHIKV nsP1 Apo i.e., native protein, nsP1 - SAM or CAPE or HC (B) DENV 3 NS5 MTase Apo i.e., native, NS5 MTase – GTP or HC or CAPE

**Supplementary Figure 4:**
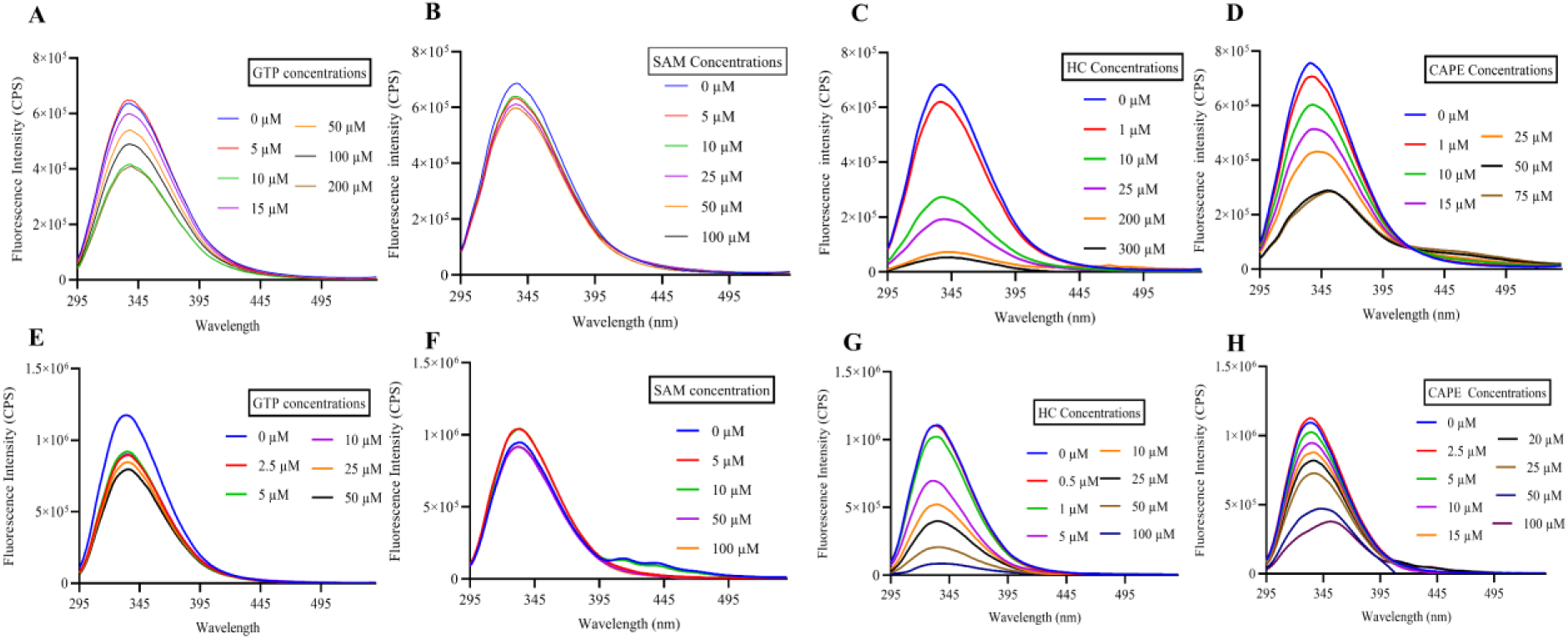
Intrinsic fluorescence intensity change in protein-compound interactions by TFS. nsP1 - (A)GTP (B) SAM (C) HC (D) CAPE. NS5 MTase – (E)GTP (D) SAM (G) HC (H) CAPE .

**Supplementary Figure 5:**
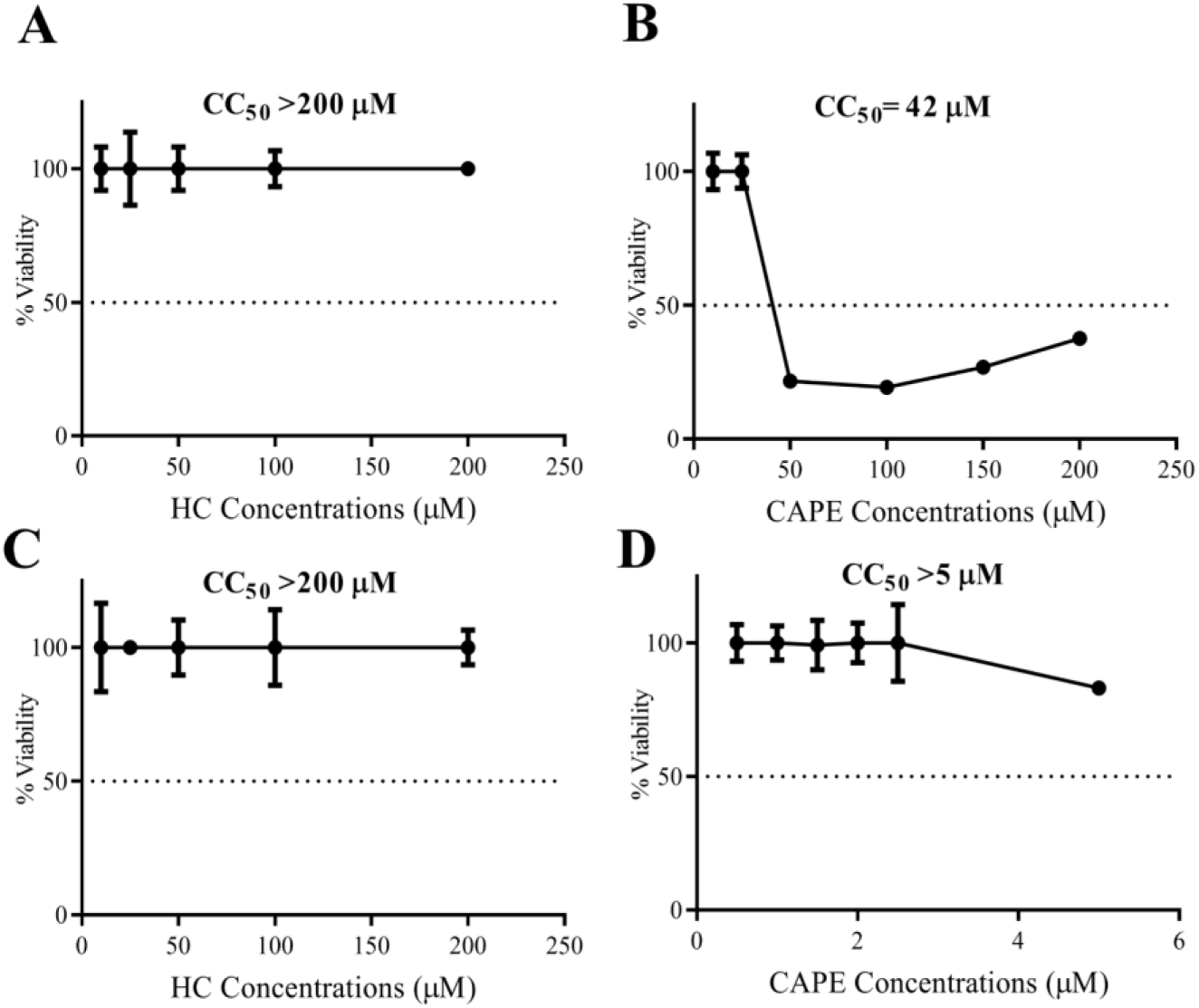
(A) and (B) depict the percent cell viability of Vero cells treated with compounds for 12 h pre-treatment followed by a 2 h incubation with 2% DMEM, and 24 h post-treatment. (C) and (D) show the percent cell viability of Vero cells treated with compounds for 12 h pre-treatment, followed by a 2 h incubation with maintenance media and 24 h post- treatment, and incubated in maintenance media for 4 days. Values are the means, and error bars represent the standard deviation from three independent experiments.The 50% cytotoxic concentration was determined based on linear dose-response analysis using GraphPad Prism 8 software. Compounds with concentrations that maintained cell viability above 90% were selected for subsequent cell culture experiments.

**Supplementary Figure 6:**
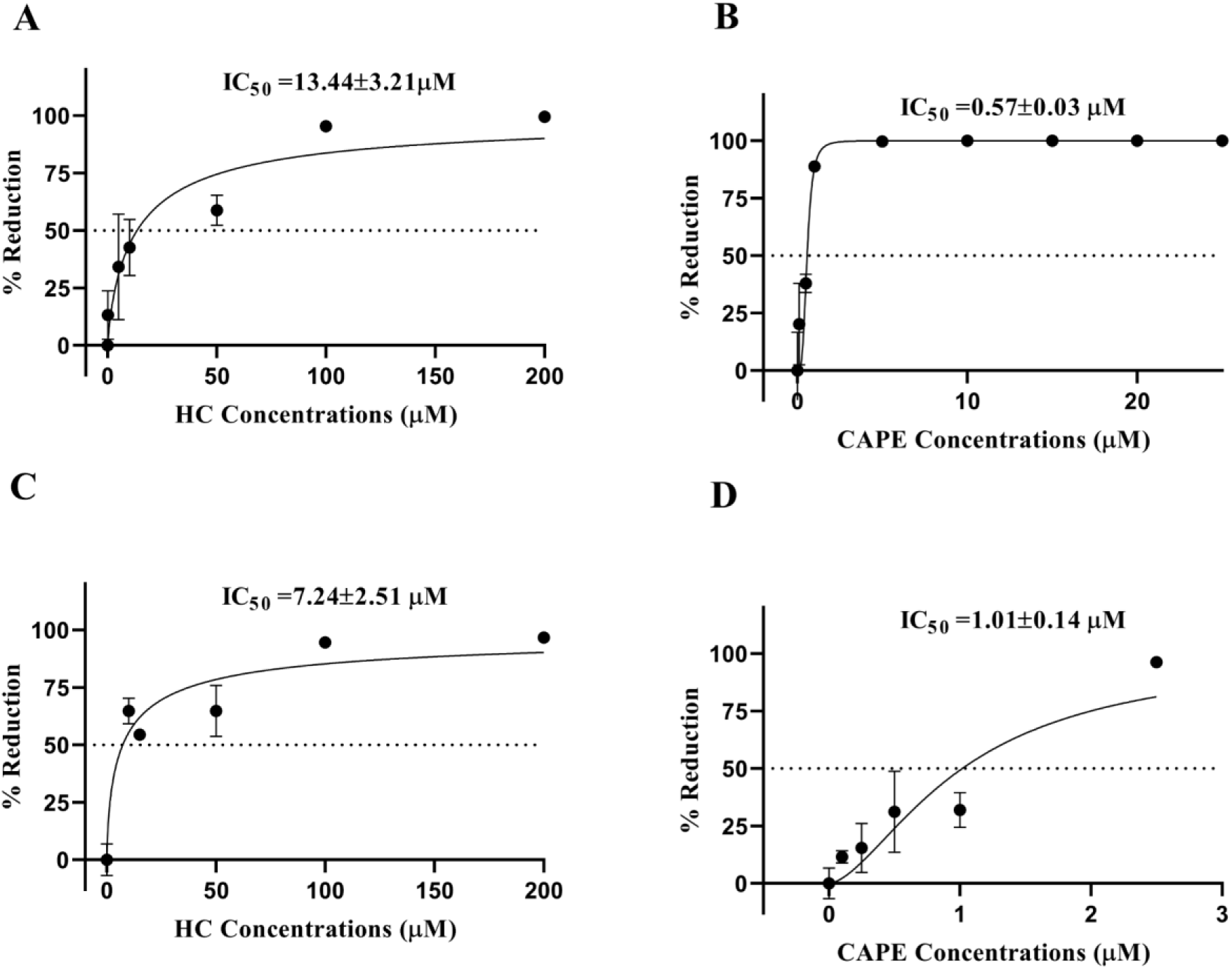
Evaluation of antiviral activity through virus titer reduction profiling: (A) HC and (B) CAPE against CHIKV; (C) HC and (D) CAPE against DENV.For this, percent inhibition was calculated using the formula: Percent Inhibition=100× ((PFU mL-1 VC−PFU mL-1 test concentration)/PFU mL-1VC), where VC represents virus control values. Data were plotted against concentration with a non-linear regression curve fit usin GraphPad Prism 8.0.

## Notes

### Competing Interest Statement

The authors have declared no competing interest.

### Summary of Updates

This version of the manuscript has been revised to include new results, which have been incorporated into the relevant sections.

